# scGNN: a novel graph neural network framework for single-cell RNA-Seq analyses

**DOI:** 10.1101/2020.08.02.233569

**Authors:** Juexin Wang, Anjun Ma, Yuzhou Chang, Jianting Gong, Yuexu Jiang, Hongjun Fu, Cankun Wang, Ren Qi, Qin Ma, Dong Xu

## Abstract

Single-cell RNA-sequencing (scRNA-Seq) is widely used to reveal the heterogeneity and dynamics of tissues, organisms, and complex diseases, but its analyses still suffer from multiple grand challenges, including the sequencing sparsity and complex differential patterns in gene expression. We introduce the scGNN (single-cell graph neural network) to provide a hypothesis-free deep learning framework for scRNA-Seq analyses. This framework formulates and aggregates cell-cell relationships with graph neural networks and models heterogeneous gene expression patterns using a left-truncated mixture Gaussian model. scGNN integrates three iterative multi-modal autoencoders and outperforms existing tools for gene imputation and cell clustering on four benchmark scRNA-Seq datasets. In an Alzheimer’s disease study with 13,214 single nuclei from postmortem brain tissues, scGNN successfully illustrated disease-related neural development and the differential mechanism. scGNN provides an effective representation of gene expression and cell-cell relationships. It is also a novel and powerful framework that can be applied to scRNA-Seq analyses.

## BACKGROUND

Single-cell RNA sequencing (scRNA-seq) techniques enable transcriptome-wide gene expression measurement in individual cells, which are essential for identifying cell type clusters, inferring the arrangement of cell populations according to trajectory topologies, and highlighting somatic clonal structures while characterizing cellular heterogeneity in complex diseases^1,2^. scRNA-seq analysis for biological inference remains challenging due to its complex and undetermined data distribution, which has an extremely large volume and high rate of dropout events. Some pioneer methodologies, e.g., Phenograph^3^, MAGIC^4^, and Seurat^5^ use a k-nearest-neighbor (KNN) graph to model the relationships between cells. However, such a graph representation may over-simplify the complex cell and gene relationships of the global cell population. Recently, the emerging graph neural network (GNN) has deconvoluted node relationships in a graph through neighbor information propagation in a deep learning architecture^6^. Compared with other autoencoders used in the scRNA-Seq analysis^7–10^ for revealing an effective representation of scRNA-Seq data via recreating its own input, the unique feature of graph autoencoder is in being able to learn a low dimensional representation of the graph topology and train node relationships in a global view of the whole graph^11^.

We introduce a multi-modal framework scGNN (single-cell graph neural network) for modeling heterogeneous cell-cell relationships and their underlying complex gene expression patterns from scRNA-Seq. scGNN trains low dimensional feature vectors (i.e., embedding) to represent relationships among cells through topological abstraction based on both gene expression and transcriptional regulation information. There are three unique features in scGNN: (*i*) scGNN utilizes GNN with multi-modal autoencoders to formulate and aggregate cell-cell relationships, providing a hypothesis-free framework to derive biologically meaningful relationships. The framework does not need to assume any statistical distribution or relationships for gene expression data or dropout events. (*ii*) Cell-type-specific regulatory signals are modeled in building a cell graph, equipped with a left-truncated mixture Gaussian (LTMG) model for scRNA-Seq data^12^. This can improve the signal-to-noise ratio in terms of embedding biologically meaningful information. (*iii*) Bottom-up cell relationships are formulated from a dynamically pruned GNN cell graph. The entire graph can be represented by pooling on learned graph embedding of all nodes in the graph. The graph embedding can be used as low-dimensional features with tolerance to noises for the preservation of topological relationships in the cell graph. The derived cell-cell relationships are adopted as regularizers in the autoencoder training to recover gene expression values.

scGNN has great potential in capturing biological cell-cell relationships in terms of cell type clustering, cell trajectory inference, cell lineages formation, and cells transitioning between states. In this paper, we mainly focus on discovering its applicative power in two fundamental aspects from scRNA-Seq data, i.e., gene imputation and cell clustering. Gene imputation aims to solve the dropout issue which commonly exists in scRNA-Seq data where the expressions of a large number of active genes are marked as zeros^13–15^. The excess of zero values often needs to be recovered or handled to avoid the exaggeration of the dropout events in many downstream biological analyses and interpretations. Existing imputation methods, such as MAGIC^4^ and SAVER^16^, have an issue in generating biased estimates of gene expression and tend to induce false-positive and biased gene correlations that could possibly eliminate some meaningful biological variations^17,18^. On the other hand, many studies, including Seurat^5^ and Phenograph^3^, have explored the cell-cell relationships using raw scRNA-seq data, and built cell graphs with reduced data dimensions and detected cell clusters by applying the Louvain modularity optimization. Accurate cell-cell relationships obey the rule that cells are more homogeneous within a cell type and more heterogeneous among different cell types^19^, The scGNN model provides a global perspective in exploring cell relationships by integrating cell neighbors on the whole population.

scGNN achieves promising performance in gene imputation and cell cluster prediction on four scRNA-Seq datasets with gold-standard cell labels^20–23^, compared to seven existing imputation and four clustering tools (**Supplementary Table S1**). We believe that the superior performance in gene imputation and cell cluster prediction benefits from (*i*) our integrative autoencoder framework, which synergistically determines cell clusters based on a bottom-up integration of detailed pairwise cell-cell relationships and the convergence of predicted clusters, and (*ii*) the integration of both gene regulatory signals and cell network representations in hidden layers as regularizers of our autoencoders. To further demonstrate the power of scGNN in complex disease studies, we applied it to an Alzheimer’s disease (AD) dataset containing 13,214 single nuclei, which elucidated its application power on cell-type identification and recovering gene expression values^24^. We claim that such a GNN-based framework is powerful and flexible enough to have great potential in integrating scMulti-Omics data.

## RESULTS

### The architecture of scGNN is comprised of stacked autoencoders

The main architecture of scGNN is used to seek effective representations of cells and genes that are useful for performing different tasks in scRNA-Seq data analyses (**Figure 1** and **Supplementary Figure S1**). It has three comprehensive computational components in an iteration process, including gene regulation integration in a feature autoencoder, cell graph representation in a graph autoencoder, gene expression updating in a set of parallel cell-type-specific cluster autoencoders, as well as the final gene expression recovery in an imputation autoencoder (**Figure 1**).

**Figure 1.**
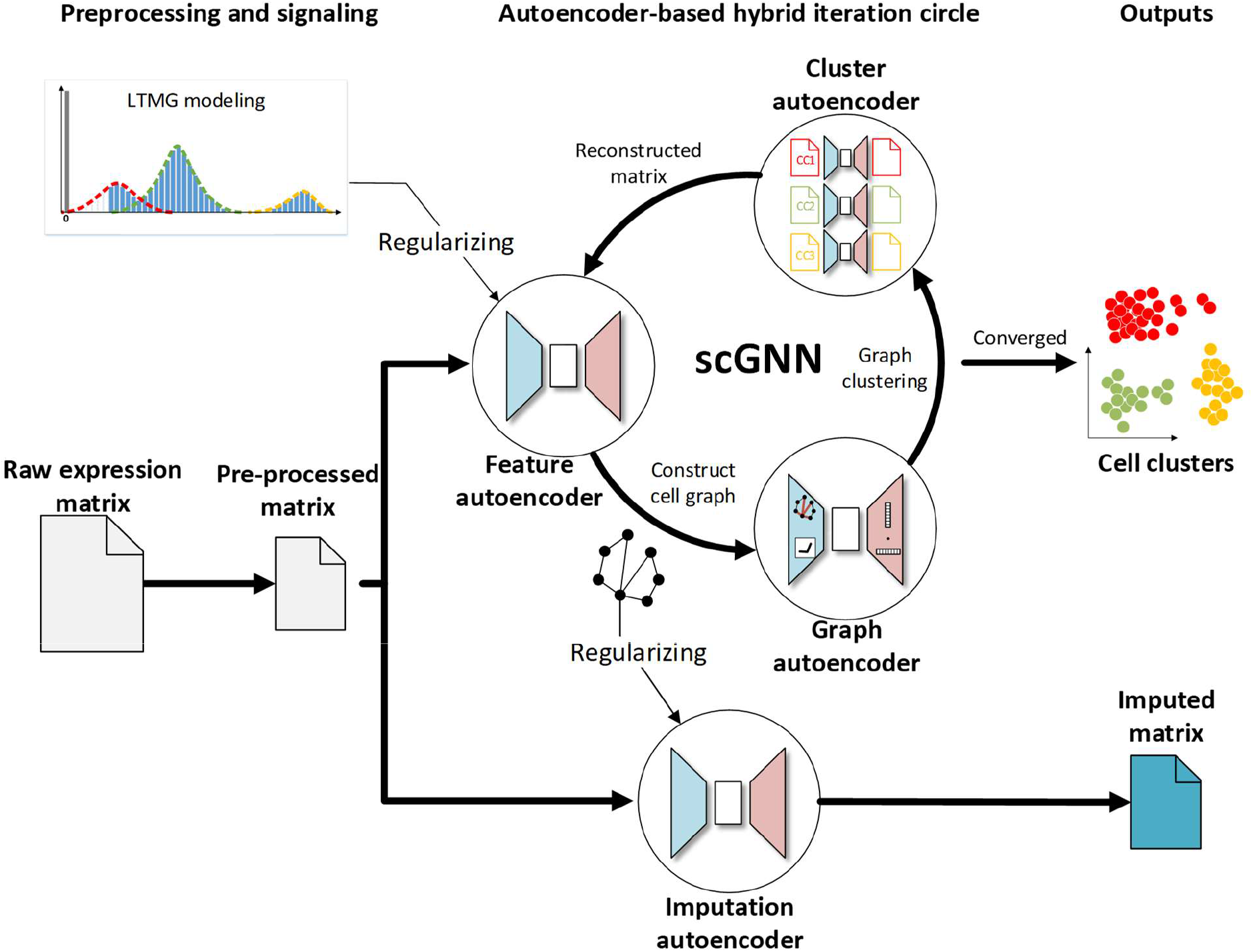
The architecture of scGNN. It takes the gene expression matrix generated from scRNA-Seq as the input. LTMG can translate the input gene expression data into a discretized regulatory signal as the regularizer for the feature autoencoder. The feature autoencoder learns a dimensional representation of the input as embedding, upon which a cell graph is constructed and pruned. The graph autoencoder learns a topological graph embedding of the cell graph, which is used for cell type clustering. The cells in each cell type have an individual cluster autoencoder to reconstruct gene expression values. The framework treats the reconstructed expression as a new input iteratively until converging. Finally, the imputed gene expression values are obtained by the feature autoencoder regularized by the cell-cell relationships in the learned cell graph on the original preprocessed raw expression matrix through the imputation autoencoder.

#### Feature autoencoder

This autoencoder intakes the pre-processed gene expression matrix after the removal of low-quality cells and genes, normalization, and variable gene ranking. First, the LTMG model^12,25^ is adopted to the top 2,000 variable genes to quantify gene regulatory signals encoded among diverse cell states in scRNA-Seq data (**Online Methods** and **Supplementary Figure S2**). This model was built based on the kinetic relationships between the transcriptional regulatory inputs and mRNA metabolism and abundance, which can infer the expression multi-modalities across single cells. The captured signals have a better signal-to-noise ratio to be used as a high-order restraint to regularize the feature autoencoder. The aim of this regularization is to treat each gene differently based on their individual regulation status through a penalty in the loss function. The feature autoencoder learns a low dimensional embedding by the gene expression reconstruction together with the regularization. A cell-cell graph is generated from the learned embedding via the KNN graph, where nodes represent individual cells and the edges represent neighborhood relations among these cells^26,27^. Then, the cell graph is pruned from selecting an adaptive number of neighbors for each node on the KNN graph by removing the noisy edges^3^.

#### Graph autoencoder

Taking the pruned cell graph as input, the encoder of the graph autoencoder uses GNN to learn a low dimensional embedding of each node and then regenerates the whole graph structure through the decoder of the graph autoencoder (**Figure 2A**). Based on the topological properties of the cell graph, the graph autoencoder abstracts intrinsic high-order cell-cell relationships propagated on the global graph. The low dimensional graph embedding integrates the essential pairwise cell-cell relationships and the global cell-cell graph topology using a graph formulation by regenerating the topological structure of the input cell graph. Then the k-means clustering method is used to cluster cells on the learned graph embedding^28^, where the number of clusters is determined by the Louvain algorithm on the cell graph.

**Figure 2.**
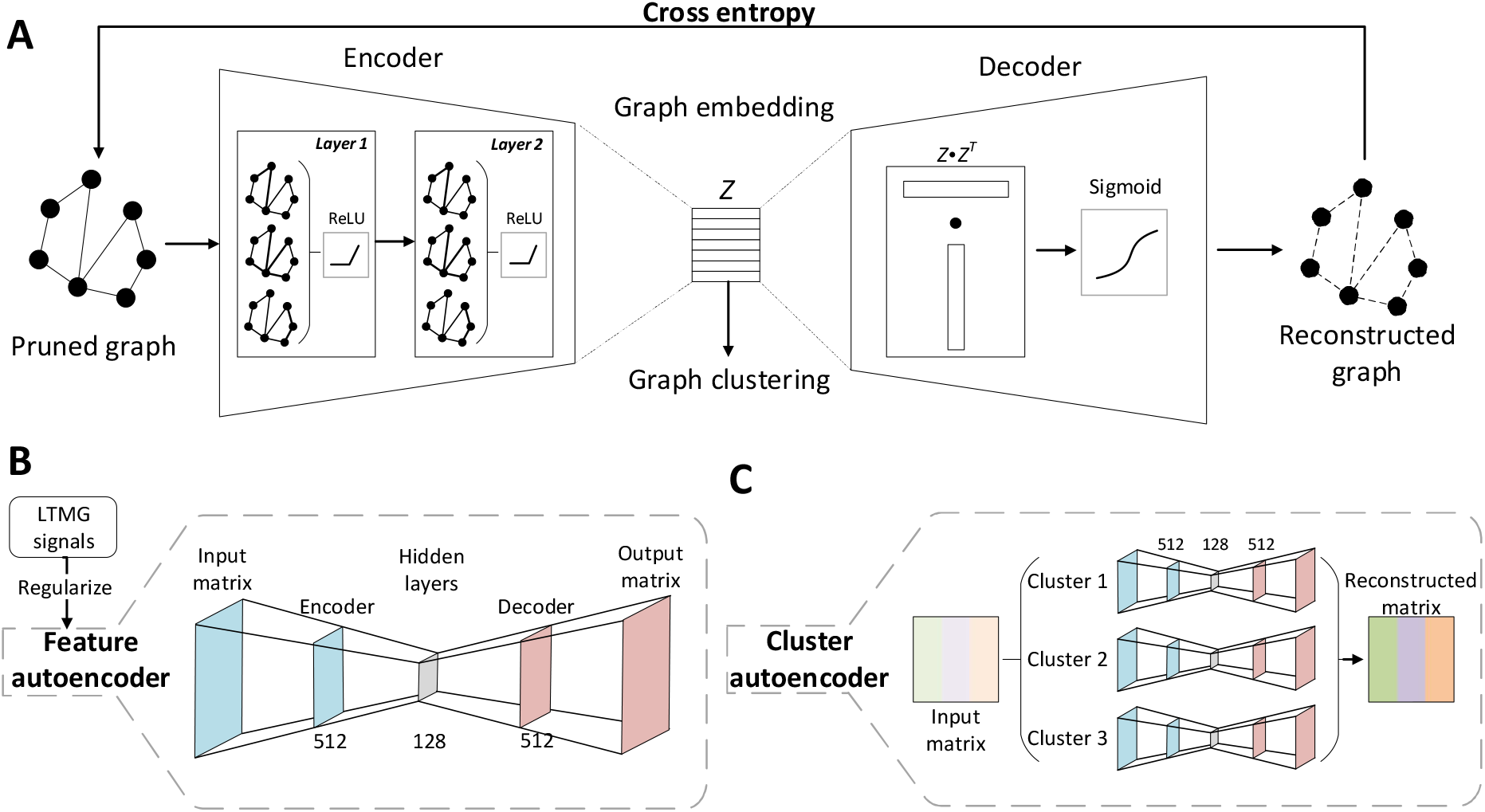
The architecture of scGNN Autoencoders. (A) The graph autoencoder takes the adjacent matrix of the pruned graph as the input. The encoder consists of two layers of GNNs. In each layer, each node of the graph aggregates information from its neighbors. The encoder learns a low dimensional presentation (i.e., graph embedding) of the pruned cell graph. The decoder reconstructs the adjacent matrix of the graph by dot products of the learned graph embedding followed by a sigmoid activation function. The graph autoencoder is trained by minimizing the cross-entropy loss between the input and the reconstructed graph. Cell clusters are obtained by applying k-means and Louvain on the graph embedding. (B) The feature autoencoder takes the expression matrix as the input, regularized by LTMG signals. The dimensions of the encoder and decoder layers are 512×128 and 128×512, respectively. The feature autoencoder is trained by minimizing the difference between the input matrix and the output matrix. (C) The cluster autoencoder takes a reconstructed expression matrix from the feature autoencoder as the input. An individual encoder is built on the cells in each of the identified clusters, and each autoencoder is trained individually. The concatenation of the results from all clusters is treated as the reconstructed matrix.

#### Cluster autoencoder

The expression matrix in each cell cluster from the feature autoencoder is reconstructed through the cluster autoencoder. Using the inferred cell type information from the graph autoencoder, the cluster autoencoder treats different cell types specifically and regenerates expression in the same cell cluster. The cluster autoencoder helps discover cell-type-specific information for each cell type in its individualized learning. Accompanied by the feature autoencoder, the cluster autoencoder leverages the inferences between global and cell-type-specific representation learning. Iteratively, the reconstructed matrix is fed back into the feature autoencoder. The iteration process stops until it converges with no change in cell clustering and this cell clustering result is recognized as the final results of cell type prediction.

#### Imputation autoencoder

After the iteration stops, this imputation autoencoder takes the original gene expression matrix as input and is trained with the additional L1 regularizer of the inferred cell-cell relationships. The regularizers (see **Online Methods**) are generated based on edges in the learned cell graph in the last iteration and their co-occurrences in the same predicted cell type. Besides, the L1 penalty term is applied to increase the model generalization by squeezing more zeroes into the autoencoder model weights. The sparsity brought by the L1 term benefits the expression imputation in dropout effects. Finally, the reconstructed gene expression values are used as the final imputation output.

### scGNN can effectively impute scRNA-Seq data and accurately predict cell clusters

To assess the imputation and cell clustering performance of scGNN, four scRNA datasets (i.e., Chung^23^, Kolodziejczy^20^, Klein^21^, and Zeisel^22^) with gold-standard cell type labels are chosen as the benchmarks (more performance evaluation on other datasets can be found in **Supplementary Materials**). We manually simulated the dropout effects by randomly flipping 10% of the non-zero entries to zeros. The median L1 distance between the original dataset and the imputed values for these corrupted entries were evaluated to compare scGNN with MAGIC^4^, SAUCIE^8^, SAVER^16^, scImpute^29^, scVI^30^, DCA^9^, and DeepImpute^31^. scGNN shows the lowest L1 distance and the highest cosine similarity in recovering leave-out values, indicating that it can accurately capture and restore true expression values (**Online Methods** and **Figure 3A**). Furthermore, scGNN depicts the underlying gene-gene relationships missed due to the sparsity of scRNA-Seq. For example, two pluripotency epiblast gene pairs, *Ccnd3* versus *Pou5f1* and *Nanog* versus *Trim28*, are lowly correlated in the original raw data but show strong positive correlations, which are differentiated by time points after scGNN imputation and, therefore, perform with a consistency leading to the desired results sought in the original paper^21^ (**Figure 3B**). The relationships of four more gene pairs are also enhanced (**Supplementary Figure S3**). In the Zeisel dataset, scGNN amplifies differentially expressed genes (DEGs) signals with a higher fold change than the original, using an imputed matrix to confidently depict the cluster heterogeneity (**Figure 3C** and **Supplementary Figure S4**).

**Figure 3.**
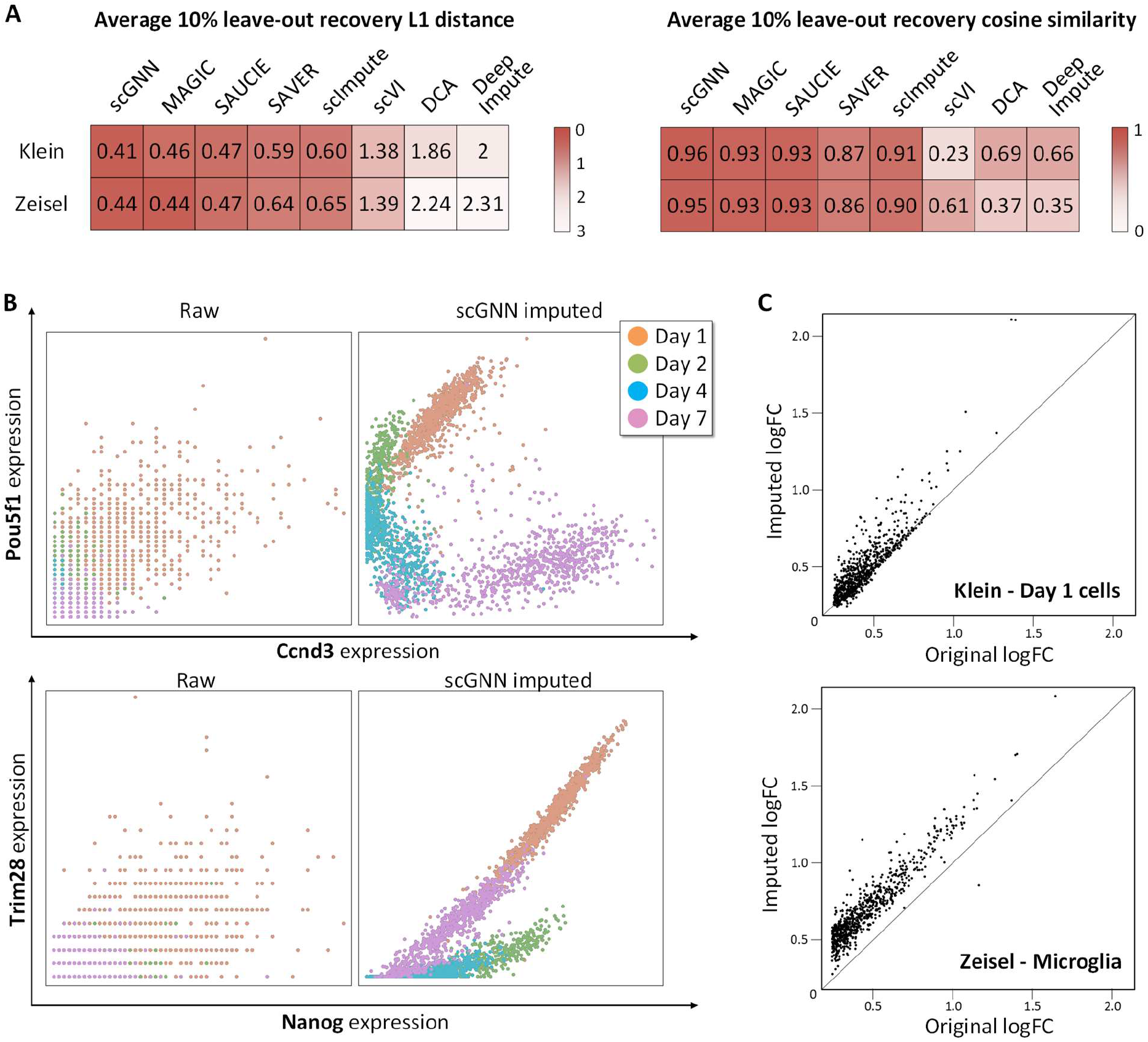
Comparison of the imputation performance. (A) The L1 distance (the lower the better) and cosine similarity (the higher the better) comparing a 10% leave-out test between scGNN and seven imputation tools on the Klein and Zeisel datasets. scGNN achieved the best scores in both datasets, indicating its superior performance in gene expression recovery. (B) Co-expression patterns can be addressed more explicitly after applying scGNN on the Klein data. No clear gene pair relationship of *Ccnd3* versus *Pou5f1* (upper panel) and *Nanog* versus *Trim28* (lower panel) is observed in the raw data (left) compared to the observation of unambiguous correlations within each cell type after scGNN imputation (right). (C) Comparison of DEG logFC scores using the original expression value (x-axis) and the scGNN imputed expression values (y-axis) identified in Day 1 cells of the Klein data (up) and Microglia cells of the Zeisel data (bottom). The differentiation signals are amplified after imputation.

Besides the artificial dropout benchmarks, we continued to evaluate the clustering performance of scGNN and the seven imputation tools on the same two datasets. The predicted cell labels were systematically evaluated using 10 criteria including an adjusted Rand index (ARI)^32^, Silhouette^33^, and eight other criteria (**Figure 4A**). By visualizing cell clustering results on UMAPs, one can observe more apparent closeness of cells within the same cluster and separation among different clusters when using scGNN embeddings compared to the other seven imputation tools (**Figure 4B**). The expression patterns show heterogeneity along with embryonic stem cell development. In the case of Klein’s time-series data, scGNN recovered a complex structure that was not well represented by the raw data, showing a well-aligned trajectory path of cell development from Day 1 to Day 7 (**Figure 4C**). Moreover, scGNN showed significant enhancement in cell clustering compared to the clustering tool (e.g., Seurat) when using the raw data (**Supplementary Figure S5**). On top of that, to address the significance of using the graph autoencoder and cluster autoencoder in scGNN, we performed ablation tests to bypass each autoencoder and compare the ARI results on the Klein dataset (**Figure 4D**). The results showed that removing either of these two autoencoders dramatically decreased the performance of scGNN in terms of cell clustering accuracy. Another test using all genes rather than the top 2,000 variable genes also showed poor performance in the results and doubled the runtime of scGNN, indicating that those low variable genes may reduce the signal-to-noise ratio and negatively affect the accuracy of scGNN. The design and comprehensive results of the ablation studies on both clustering and imputation are detailed in **Supplementary Method** and **Table S2-S7** and **S11**. We also extensively studied the parameter selection in **Supplementary Table S8-S10** and **S12**.

**Figure 4.**
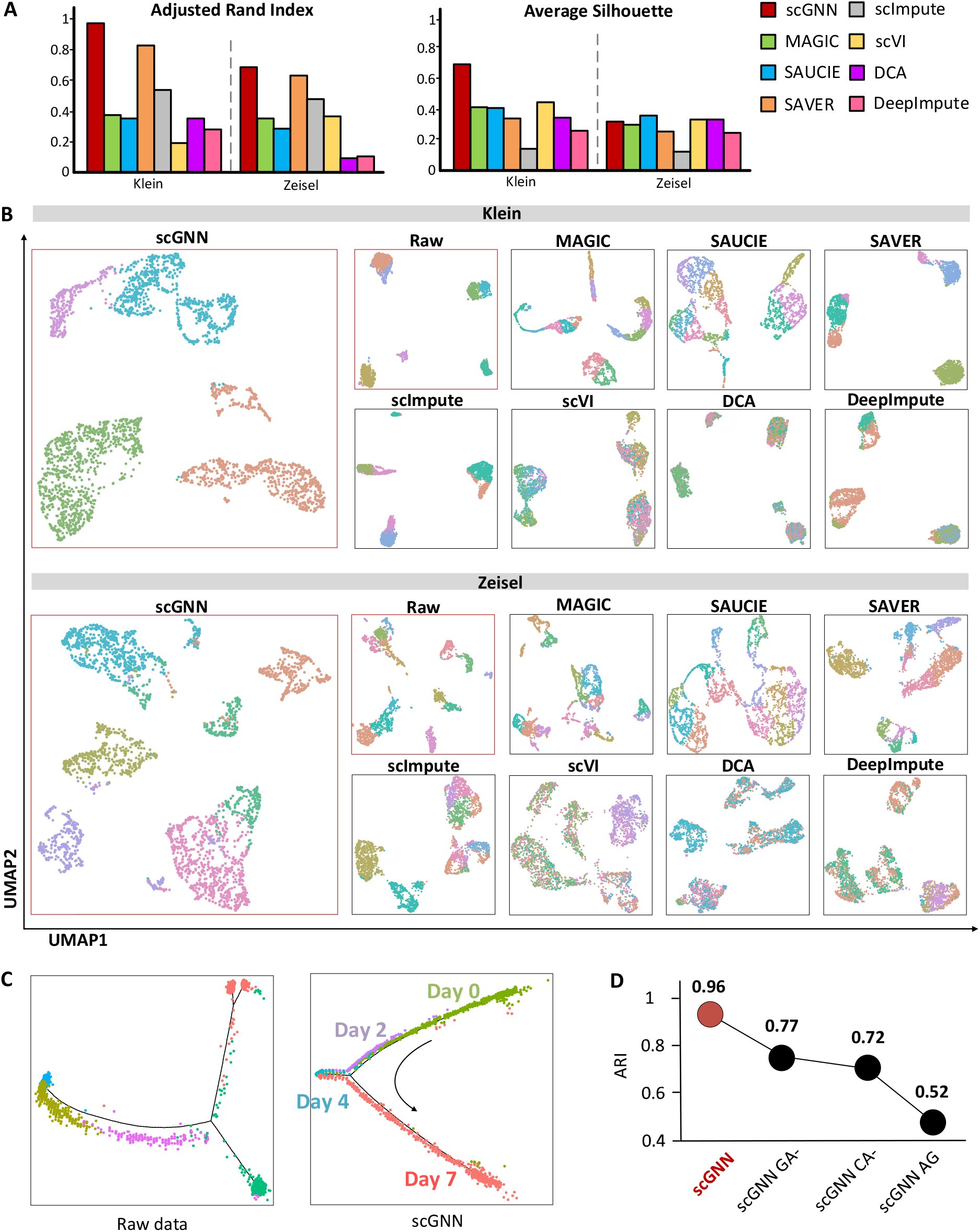
Cell clustering and trajectory evaluations. (A) Comparison of ARI and Silhouette scores among scGNN and seven tools using Klein and Zeisel datasets. (B) Comparison of UMAP visualizations on the same two datasets, indicating that when scGNN embeddings are utilized, cells are more closely grouped within the same cluster but when other tools are used, cells are more separated between clusters. Raw data is clustered and visualized using Seurat. (C) Pseudotime analysis using the raw expression matrix and scGNN imputed matrix of the Klein dataset via Monocle2. (D) Justification of using the graph autoencoder, the cluster autoencoder, and the top 2,000 variable genes on the Klein dataset in the scGNN framework, in terms of ARI. scGNN CA-shows the results of the graph autoencoder’s ablation, CA-shows the results of the cluster autoencoder’s ablation, and AG shows the results after using all genes in the framework.

### scGNN illustrates AD-related neural development and the underlying regulatory mechanism

To further demonstrate the applicative power of scGNN, we applied it to a scRNA-Seq dataset (GEO accession number GSE138852) containing 13,214 single nuclei collected from six AD and six control brains^34^. scGNN identifies 10 cell clusters, including microglia, neurons, oligodendrocyte progenitor cells (OPCs), astrocytes, and six sub-clusters of oligodendrocytes (**Figure 5A**). Specifically, the proportions of these six oligodendrocyte sub-clusters differ between AD patients (Oligos 2, 3, and 4) and healthy controls (Oligos 1, 5, and 6) (**Figure 5B**). Moreover, the difference between AD and the control in the proportion of astrocyte and OPCs is observed, indicating the change of cell population in AD patients compared to healthy controls (**Figure 5B**). We then combined these six oligodendrocyte sub-clusters into one to discover DEGs. Since scGNN can significantly increase true signals in the raw dataset, DEG patterns are more explicit (**Supplementary Figure S6**). Among all DEGs, we confirmed 22 genes as cell-type-specific markers for astrocytes, OPCs, oligodendrocytes, and neurons, in that order^35^ (**Figure 5C**). A biological pathway enrichment analysis shows several highly positive-enrichments in AD cells compared to control cells among all five cell types. These enrichments include oxidative phosphorylation and pathways associated with AD, Parkinson’s disease, and Huntington disease^36^ (**Figure 5D** and **Supplementary Figure S7**). Interestingly, we observed a strong negative enrichment of the MAPK (mitogen-activated protein kinase) signaling pathway in the microglia cells, suggesting a relatively low MAPK regulation in microglia than other cells.

**Figure 5.**
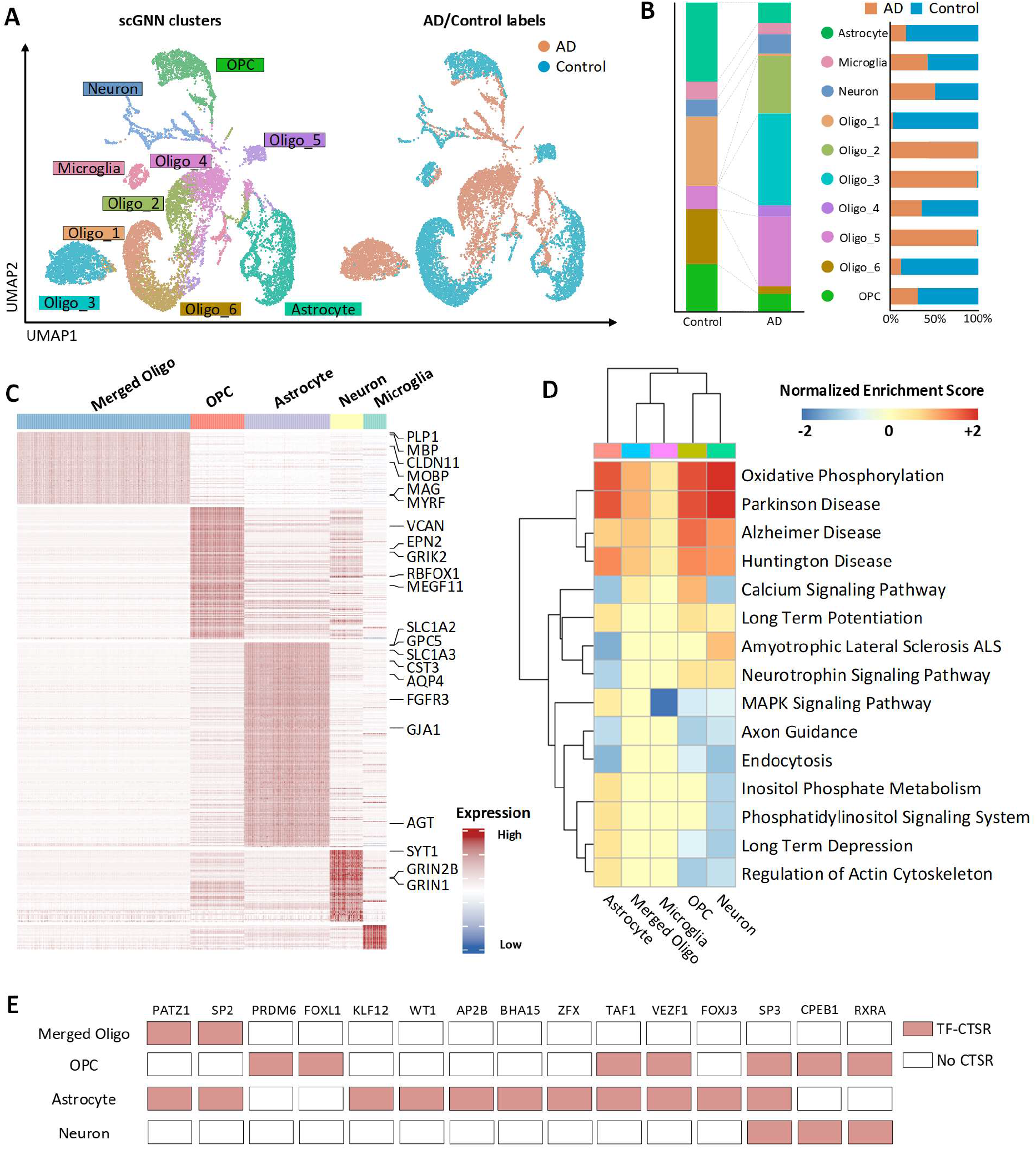
Alzheimer’s disease dataset (GSE138852) analysis based on scGNN. (A) Cell clustering UMAP. Labeled with scGNN clusters (left) and AD/control samples (right). (B) Comparison of cell proportions in AD/control samples (left) and each cluster (right). (C) Heatmap of DEGs (logFC > 0.25) in each cluster. Six oligodendrocyte sub-clusters are merged as one to compare with other cell types. Marker genes identified in DEGs are listed on the right. (D) Selected AD-related enrichment pathways in each cell type in the comparison between AD and control cells. (E) Underlying TFs are responsible for the cell-type-specific gene regulations identified by IRIS3.

In order to investigate the regulatory mechanisms underlying the AD-related neural development, we applied the imputed matrix of scGNN to IRIS3 (an integrated cell-type-specific regulon inference server from single-cell RNA-Seq) and identified 21 cell-type-specific regulons (CTSR) in five cell types^37^ (**Figure 5E** and **Supplementary Table S13**; IRIS3 job ID: 20200626160833). Not surprisingly, we identified several AD-related transcription factors (TFs) and target genes that have been reported to be involved in the development of AD. SP2 is a common TF identified in both oligodendrocytes and astrocytes. It has been shown to regulate the *ABCA7* gene, which is an IGAP (International Genomics of Alzheimer’s Project) gene that is highly associated with late-onset AD^38^. We also observed an SP2 CTSR in astrocytes that regulate *APOE, AQP4, SLC1A2, GJA1, and FGFR3*. All of these five targeted genes are marker genes of astrocytes, which have been reported to be associated with AD^39,40^. In addition, the SP3 TF is identified in all cell clusters which can regulate the synaptic function in neurons, and it is extremely activated in AD^41,42^. We identified CTSRs regulated by SP3 in OPCs, astrocytes, and neurons suggesting a significant SP3 related regulation shifts in these three clusters. We observed 26, 60, and 22 genes that were uniquely regulated in OPCs, astrocytes, and neurons, as well as 60 genes shared among the three clusters (**Supplementary Table S14**). Such findings provide a direction for the discovery of SP3 function in AD studies.

## DISCUSSION

It is still a fundamental challenge to explore cellular heterogeneity in high-volume, high-sparsity, and noisy scRNA-Seq data, where the high-order topological relationships of the whole-cell graph are still not well explored and formulated. The key innovations of scGNN are incorporating global propagated topological features of the cells through GNNs, together with integrating gene regulatory signals in an iterative process for scRNA-Seq data analysis. The benefits of GNN is its intrinsic learnable properties of propagating and aggregating attributes to capture relationships across the whole cell-cell graph. Hence, the learned graph embedding can be treated as the high-order representations of cell-cell relationships in scRNA-Seq data in the context of graph topology. Unlike the previous autoencoder applications in scRNA-Seq data analysis, which only captures the top-down distributions of the overall cells, scGNN can effectively aggregate detailed relationships between similar cells using a bottom-up approach. Furthermore, scGNN integrates gene regulatory signals efficiently by representing them discretely in LTMG in the feature autoencoder regularization. These gene regulatory signals can help identify biologically meaningful gene-gene relationships as they apply to our framework and eventually, they are proven capable of enhancing performance. Technically, scGNN adopts multi-modal autoencoders in an iterative manner to recover gene expression values and cell type prediction simultaneously. Notably, scGNN is a hypothesis-free deep learning framework on a data-driven cell graph model, and it is flexible to incorporate different statistical models (e.g. LTMG) to analyze complex scRNA-Seq datasets.

Some limitations can still be found in scGNN. (*i*) It is prone to achieve better results with large datasets, compared to relatively small datasets (e.g., less than 1,000 cells), as it is designed to learn better representations with many cells from scRNA-Seq data, as shown in the benchmark results, and (*ii*) Compared with statistics model-based methods, the iterative autoencoder framework needs more computational resources, which is more time-consuming and less interpretable. In the future, we will investigate creating a more efficient scGNN model with a lighter and more compressed architecture.

In the future, we will continue to enhance scGNN by implementing heterogeneous graphs to support the integration of single-cell multi-omics data (e.g., the intra-modality of Smart-Seq2 and Droplet scRNA-Seq data; and the inter-modality integration of scRNA-Seq and scATAC-Seq data). We will also incorporate attention mechanisms and graph transformer models^43^ to make the analyses more explainable. Specifically, by allowing the integration of scRNA-Seq and scATAC-Seq data, scGNN has the potential to elucidate cell-type-specific gene regulatory mechanisms^44^. On the other hand, T cell receptor repertoires are considered as unique identifiers of T cell ancestries that can improve both the accuracy and robustness of predictions regarding cell-cell interactions^45^. scGNN can also facilitate batch effects and build connections across diverse sequencing technologies, experiments, and modalities. Moreover, scGNN can be applied to analyze spatial transcription datasets regarding spatial coordinates as additional regularizers to infer the cell neighborhood representation and better prune the cell graph. We plan to develop a more user-friendly software system from our scGNN model, together with modularized analytical functions in support of standardizing the data format, quality control, data integration, multi-functional scMulti-seq analyses, performance evaluations, and interactive visualizations.

## ONLINE METHODS

### Dataset preprocessing

scGNN takes the scRNA-Seq gene expression profile as the input. Data filtering and quality control are the first steps of data preprocessing. Due to the high dropout rate of scRNA-seq expression data, only genes expressed as nonzero in more than 1% of cells, and cells expressed as nonzero in more than 1% of genes are kept. Then, genes are ranked by standard deviation, i.e., the top 2,000 genes in variances are used for the study. All the data are log-transformed.

### Left Truncated Mixed Gaussian (LTMG) modeling

A mixed Gaussian model with left truncation assumption is used to explore the regulatory signals from gene expression^12^. The normalized expression values of gene *X* over *N* cells are denoted as *X* = {*x*_1_, …*x*_*N*_}, where *x*_*j*_ *∈ X* is assumed to follow a mixture of *k* Gaussian distributions, corresponding to *k* possible gene regulatory signals (**TRSs**). The density function of *X* is:

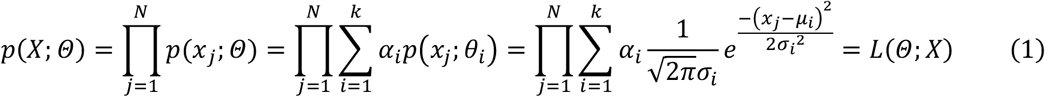

where *α*_*i*_ is the mixing weight, *μ*_*i*_ and *σ*_*i*_ are the mean and standard deviation of the *i^th^* Gaussian distribution, which can be estimated by: 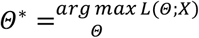 to model the errors at zero and the low expression values. With the left truncation assumption, the gene expression profile is split into *M*, which is a truly measured expression of values, and *N* − *M* representing left-censored gene expressions for *N* conditions. The parameter *Θ* maximizes the likelihood function and can be estimated by an expectation-maximization algorithm. The number of Gaussian components is selected by the Bayesian Information Criterion; then, the original gene expression values are labeled to the most likely distribution under each cell. In detail, the probability that *x*_*j*_ belongs to distribution *i* is formulated by:

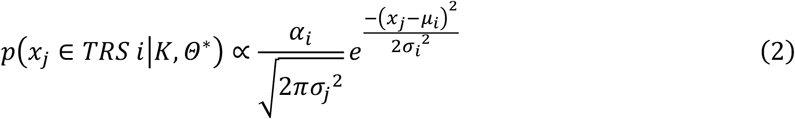

where *x*_*j*_ is labeled by TRS *i* if 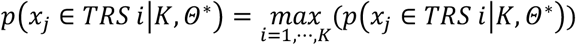. Thus, the discrete values (1,2, …, *K*) for each gene are generated.

### Feature autoencoder

The feature autoencoder is proposed to learn the representative embedding of the scRNA expression through stacked two layers of dense networks in both the encoder and decoder. The encoder constructs the low dimensional embedding of *X*’ from the input gene expression *X*, and the encoder reconstructs the expression 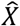 from the embedding; thus, 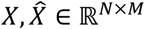 and *X*’∈ ℝ^*N×M′*^, where *M* is the number of input genes, *M*′ is the dimension of the learned embedding, and *M*′ < *M*. The objective of training the feature autoencoder is to achieve a maximum similarity between the original and reconstructed through minimizing the loss function, in which 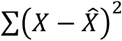 is the main term serving as the mean squared error (**MSE**) between the original and the reconstructed expressions.

### Regularization

Regularization is adopted to integrate gene regulation information during the feature autoencoder training process. The aim of this regularization is to treat each gene differently based on their individual gene regulation role through penalizing it in the loss function. In each cell, the MSE of each gene is element-wise multiplication with discrete gene regulation signals from TRS, as defined in *Eq.(5)*.

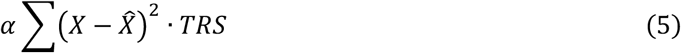

where *α* is a parameter used to control the strength of gene regulation regularization; *α* ∈ [0,1]. Thus, the loss function of the feature autoencoder is shown as *Eq.(6)*.

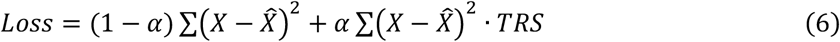

In the encoder, the output dimensions of the first and second layers are set as 512 and 128, respectively. Each layer is followed by the ReLU activation function. In the decoder, the output dimensions of the first and second layers are 128 and 512, respectively. Each layer is followed by a sigmoid activation function. The learning rate is set as 0.001. The cluster autoencoder has the same architecture as the feature autoencoder, but without gene regulation regularization in the loss function.

### Cell graph and pruning

The cell graph formulates the cell-cell relationships using embedding learned from the feature autoencoder. As done in previous works^4,46^, the cell graph is built from a KNN graph, where nodes are individual single-cells, and the edges are relationships between cells. *K* is the predefined parameter used to control the scale of the captured interaction between cells. Each node finds its neighbors within the *K* shortest distances and creates edges between them and itself. Euclidian distance is calculated as the weights of the edges on the learned embedding vectors. The pruning process selects an adaptive number of neighbors for each node on the original KNN graph and keeps a more biologically meaningful cell graph. Here, Isolation Forest is applied to prune the graph to detect the outliner in the *K*-neighbors of each node^47^. Isolation Forest builds individual random forest to check distances from the node to all *K* neighbors and only disconnects the outliners.

### Graph autoencoder

The graph autoencoder learns to embed and represent the topological information from the pruned cell graph. For the input pruned cell graph, *G* = (*V*, *E*) with *N* = |*V*| nodes denoting the cells and *E* representing the edges. *A* is its adjacency matrix and *D* is its degree matrix. The node feature matrix of the graph autoencoder is the learned embedding *X*′ from the feature autoencoder.

The graph convolution network (GCN) is defined as 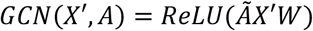, and *W* is a weight matrix learned from the training. 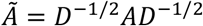 is the symmetrically normalized adjacency matrix and activation function *ReLU*(∙) = *max* (0,∙). The encoder of the graph autoencoder is composed of two layers of GCN, and *Z* is the graph embedding learned through the encoder in *Eq.(7)*. *W*_1_ and *W*_2_ are learned weight matrices in the first and second layers, and the output dimensions of the first and second layers are set at 32 and 16, respectively. The learning rate is set at 0.001.

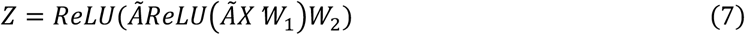

The decoder of the graph autoencoder is defined as an inner product between the embedding:

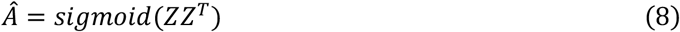

where 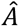 is the reconstructed adjacent matrix of *A. sigmoid(∙)* = 1/(1 + *e*^−∙^) is the sigmoid activation function.

The goal of learning the graph autoencoder is to minimize the cross-entropy *L* between the input adjacent matrix *A* and the reconstructed matrix 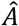.

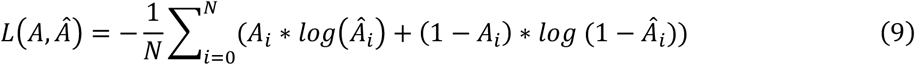

where *A*_*i*_ and 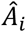 are the elements of adjacent matrix *A* and 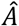. *N* is the total number of elements in the adjacent matrix.

### Iterative process

The iterative process aims to build the single-cell graph iteratively until converging. The iterative process of the cell graph can be defined as:

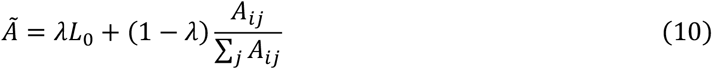

where *L*_0_ is the normalized adjacency matrix of the initial pruned graph, and 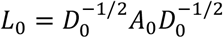, where *D*_0_ is the degree matrix. λ is the parameter to control the converging speed, λ ∈ [0,1]. Each time in iteration *t*, two criteria are checked to determine whether to stop the iteration: (1) that is, to determine whether the adjacency matrix converges, i.e., 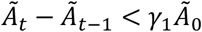, or (2) whether the inferred cell types are similar enough, i.e., *ARI* < *γ*_2_. ARI is the similarity measurement, which is detailed in the next section. In our setting, λ = 0.5 and *γ*_1_, *γ*_2_ = 0.99. The cell type clustering results obtained in the last iteration are chosen as the final cell type results.

### Imputation autoencoder

After the iterative process stops, the imputation autoencoder imputes and denoises the raw expression matrix within the inferred cell-cell relationship. The imputation autoencoder shares the same architecture as the feature autoencoder, but it also uses three additional regularizers from the cell graph in *Eq*.(11), cell types in *Eq*.(12), and the L1 regularizer in *Eq*.(13).

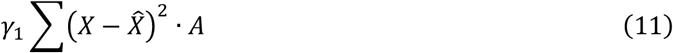

where *A* is the adjacent matrix from the pruned cell graph in the last iteration. Cells within an edge in the pruned graph will be penalized in the training.

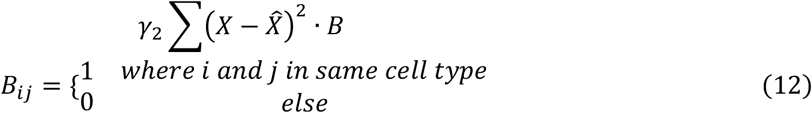

where *B* is the relationship matrix between cells, and two cells in the same cell type have a *B_ij_* value of 1. Cells within the same inferred cell type will be penalized in the training. *γ*_1_,*γ*_2_ are the intensities of the regularizers and *γ*_1_, *γ*_2_ ∈ [0,1]. The L1 regularizer is defined as

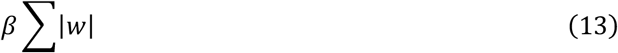

which brings sparsity and increases the generalization performance of the autoencoder by reducing the number of non-zero *w* terms in ∑|*w*|, where *β* is a hyper-parameter controlling the intensity of the L1 term (*β* ∈ [0,1]). Therefore, the loss function of the imputation autoencoder is

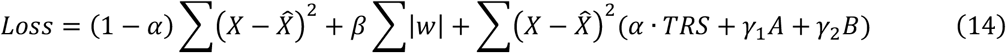

### Benchmark evaluation compared to existing tools

#### Imputation evaluation

For benchmarking imputation performance, we added noises by randomly flipping 10% of the nonzero entries to zero to mimic the dropout effects. We evaluated both the median L1 distance and cosine similarity between the original dataset and the imputed values for these corrupted entries. For all the flipped entries, *x* is the row vector of the original expression, and *y* is its corresponding row vector of the imputed expression. The L1 distance is the absolute deviation between the value of the original and imputed expression. A lower L1 distance means a higher similarity.

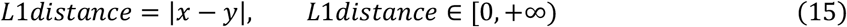

The cosine similarity computes the dot products between original and imputed expression.

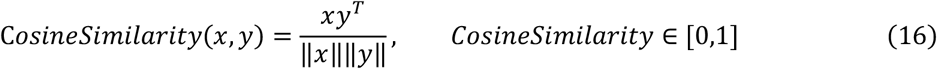

The process is repeated three times, and the mean and standard deviation were selected as a comparison. The scores are compared between scGNN and seven imputation tools (i.e., MAGIC^4^, SAUCIE^8^, SAVER^16^, scImpute^29^, scVI^30^, DCA^9^, and DeepImpute^31^), all using the default parameters.

#### Clustering evaluation

We compared the cell clustering results of scGNN, the same seven imputation tools, and four clustering tools (i.e., Seurat^5^, CIDR^48^, Monocle^49^, and RaceID^50^), in terms of ten clustering evaluation scores. The default parameters are applied in all test tools. ARI^32^ is used to compute similarities by considering all pairs of the samples that are assigned in clusters in the current and previous clustering adjusted by random permutation.

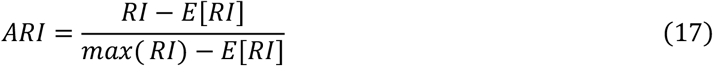

where the unadjusted rand index (RI) is defined as

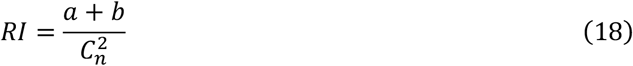

where *a* is the number of pairs correctly labeled in the same sets, and b is the number of pairs correctly labeled as not in the same dataset. 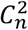 is the total number of possible pairs. *E*[*RI*] is the expected RI of random labeling. More quantitative measurements are also used in the **Supplemental Materials**.

### Case study of the AD database

We applied scGNN on a public Alzheimer’s disease (AD) scRNA-Seq data with 13,214 cells^24^. The resolution of scGNN was set to 1.0, *K*I was set to 20, and the remaining parameters were kept as default. The AD patient and control labels were provided by the original paper and used to color the cells on the same UMAP coordinates generated from scGNN. We simply combined cells in six oligodendrocyte subpopulations into one cluster, referred to as merged oligo. The DEGs were identified in each cell cluster via the Wilcoxon rank-sum test implemented in the Seurat package along with adjusted p-values using the Benjamini-Hochberg procedure with a nominal level of 0.05. DEGs with logFC > 0.25 or < − 0.25 were finally selected. We further identified the DEGs between AD and control cells in each cluster using the same strategy and applied GSEA for pathway enrichment analysis^51^. The imputed matrix, which resulted from scGNN was then sent to IRIS3 for CTSR prediction, using the predicted cell clustering labels with merged oligodendrocytes^37^. The default parameters were served in regulatory analysis in IRIS3.

## Supporting information

Supplementary Figure S1-S7, Table S1-S2, Methods S1-S4, Algorithm S1

## Data availability

Three benchmark and AD case datasets can be downloaded from GEO databases with accession numbers of: GSE75688 (the Chung data); GSE65525 (the Klein data); GSE60361 (the Zeisel data); and GSE138852 (AD case). The Kolodziejczy data can be accessed from EBI with an accession number of E-MTAB-2600.

## Software Implementation

Tools and packages used in this paper include: Python version 3.7.6, numpy version 1.18.1, torch version 1.4.0, networkx version 2.4, pandas version 0.25.3, rpy2 version 3.2.4, matplotlib version 3.1.2, seaborn version 0.9.0, umap-learn version 0.3.10, munkres version 1.1.2, R version 3.6.1, and igraph version 1.2.5. The IRIS3 website is at https://bmbl.bmi.osumc.edu/iris3/index.php.

## CODE AVAILABILITY

Our tool is open source and publicly available at GitHub (https://github.com/scgnn/scGNN).

## ACKNOWLEDGEMENTS

This work was supported by the National Institutes of Health’s National Institute of General Medical Sciences (awards R35-GM126985 and R01-GM131399).

## AUTHOR CONTRIBUTIONS

Conceptualization: Q.M., D.X.; Methodology: J.W., A.M.; Software: J.W., C.Y.; Investigation: J.W., Q.R.; Formal Analysis: A.M., J.W., J.G., Y.C., J.Y. Resources and Reagents: J.W., J.G., R.Q.; Writing, Review, and Editing: J.W., A.M., H.F., Q.M., D.X.

## COMPETING INTERESTS

The authors declare no competing interests.

## Notes

### Competing Interest Statement

The authors have declared no competing interest.

## REFERENCES

1 Hwang, B., Lee, J. H. & Bang, D. Single-cell RNA sequencing technologies and bioinformatics pipelines. Exp Mol Med 50, 96, doi:10.1038/s12276-018-0071-8 (2018).

2 Gawel, D. R. et al. A validated single-cell-based strategy to identify diagnostic and therapeutic targets in complex diseases. Genome Med 11, 47, doi:10.1186/s13073-019-0657-3 (2019).

3 Levine, J. H. et al. Data-Driven Phenotypic Dissection of AML Reveals Progenitor-like Cells that Correlate with Prognosis. Cell 162, 184–197, doi:10.1016/j.cell.2015.05.047 (2015).

4 van Dijk, D. et al. Recovering Gene Interactions from Single-Cell Data Using Data Diffusion. Cell 174, 716–729 e727, doi:10.1016/j.cell.2018.05.061 (2018).

5 Butler, A., Hoffman, P., Smibert, P., Papalexi, E. & Satija, R. Integrating single-cell transcriptomic data across different conditions, technologies, and species. Nat Biotechnol 36, 411–420, doi:10.1038/nbt.4096 (2018).

6 Kipf, T. N. & Welling, M. Semi-Supervised Classification with Graph Convolutional Networks. The International Conference on Learning Representations (ICLR) (2017).

7 Wang, W., Huang, Y., Wang, Y. & Wang, L. Generalized Autoencoder: A Neural Network Framework for Dimensionality Reduction. 2014 IEEE Conference on Computer Vision and Pattern Recognition Workshops, 496–503, doi:10.1109/CVPRW.2014.79 (2014).

8 Amodio, M. et al. Exploring single-cell data with deep multitasking neural networks. Nat Methods 16, 1139–1145, doi:10.1038/s41592-019-0576-7 (2019).

9 Eraslan, G., Simon, L. M., Mircea, M., Mueller, N. S. & Theis, F. J. Single-cell RNA-seq denoising using a deep count autoencoder. Nat Commun 10, 390, doi:10.1038/s41467-018-07931-2 (2019).

10 Miao, Z. et al. Putative cell type discovery from single-cell gene expression data. Nature Methods 17, 621–628, doi:10.1038/s41592-020-0825-9 (2020).

11 Kipf, T. N. & Welling, M. Variational Graph Auto-Encoders. arXiv e-prints, arXiv:1611.07308 (2016). <https://ui.adsabs.harvard.edu/abs/2016arXiv161107308K>.

12 Wan, C. et al. LTMG: a novel statistical modeling of transcriptional expression states in single-cell RNA-Seq data. Nucleic Acids Res 47, e111, doi:10.1093/nar/gkz655 (2019).

13 Huang, M. et al. SAVER: gene expression recovery for single-cell RNA sequencing. Nature Methods 15, 539 (2018).

14 Risso, D., Perraudeau, F., Gribkova, S., Dudoit, S. & Vert, J. P. A general and flexible method for signal extraction from single-cell RNA-seq data. Nat Commun 9, 284, doi:10.1038/s41467-017-02554-5 (2018).

15 Svensson, V. et al. Power analysis of single-cell RNA-sequencing experiments. Nat Methods 14, 381–387, doi:10.1038/nmeth.4220 (2017).

16 Huang, M. et al. SAVER: gene expression recovery for single-cell RNA sequencing. Nat Methods 15, 539–542, doi:10.1038/s41592-018-0033-z (2018).

17 Wang, J. et al. Data denoising with transfer learning in single-cell transcriptomics. Nat Methods 16, 875–878, doi:10.1038/s41592-019-0537-1 (2019).

18 Zhang, L. & Zhang, S. Comparison of Computational Methods for Imputing Single-Cell RNA-Sequencing Data. IEEE/ACM Trans Comput Biol Bioinform 17, 376–389, doi:10.1109/TCBB.2018.2848633 (2020).

19 Liu, B. et al. An entropy-based metric for assessing the purity of single cell populations. Nature Communications 11, 3155, doi:10.1038/s41467-020-16904-3 (2020).

20 Kolodziejczyk, A. A. et al. Single Cell RNA-Sequencing of Pluripotent States Unlocks Modular Transcriptional Variation. Cell Stem Cell 17, 471–485, doi:10.1016/j.stem.2015.09.011 (2015).

21 Klein, A. M. et al. Droplet barcoding for single-cell transcriptomics applied to embryonic stem cells. Cell 161, 1187–1201, doi:10.1016/j.cell.2015.04.044 (2015).

22 Zeisel, A. et al. Brain structure. Cell types in the mouse cortex and hippocampus revealed by single-cell RNA-seq. Science 347, 1138–1142, doi:10.1126/science.aaa1934 (2015).

23 Chung, W. et al. Single-cell RNA-seq enables comprehensive tumour and immune cell profiling in primary breast cancer. Nat Commun 8, 15081, doi:10.1038/ncomms15081 (2017).

24 Grubman, A. et al. A single-cell atlas of entorhinal cortex from individuals with Alzheimer’s disease reveals cell-type-specific gene expression regulation. Nat Neurosci 22, 2087–2097, doi:10.1038/s41593-019-0539-4 (2019).

25 Xie, J. et al. QUBIC2: a novel and robust biclustering algorithm for analyses and interpretation of large-scale RNA-Seq data. Bioinformatics 36, 1143–1149, doi:10.1093/bioinformatics/btz692 (2020).

26 Bendall, S. C. et al. Single-cell trajectory detection uncovers progression and regulatory coordination in human B cell development. Cell 157, 714–725, doi:10.1016/j.cell.2014.04.005 (2014).

27 Wolf, F. A. et al. PAGA: graph abstraction reconciles clustering with trajectory inference through a topology preserving map of single cells. Genome Biol 20, 59, doi:10.1186/s13059-019-1663-x (2019).

28 Blondel, V. D., Guillaume, J.-L., Lambiotte, R. & Lefebvre, E. Fast unfolding of communities in large networks. Journal of Statistical Mechanics: Theory and Experiment 2008, P10008, doi:10.1088/1742-5468/2008/10/p10008 (2008).

29 Li, W. V. & Li, J. J. An accurate and robust imputation method scImpute for single-cell RNA-seq data. Nat Commun 9, 997, doi:10.1038/s41467-018-03405-7 (2018).

30 Lopez, R., Regier, J., Cole, M. B., Jordan, M. I. & Yosef, N. Deep generative modeling for single-cell transcriptomics. Nat Methods 15, 1053–1058, doi:10.1038/s41592-018-0229-2 (2018).

31 Arisdakessian, C., Poirion, O., Yunits, B., Zhu, X. & Garmire, L. X. DeepImpute: an accurate, fast, and scalable deep neural network method to impute single-cell RNA-seq data. Genome Biol 20, 211, doi:10.1186/s13059-019-1837-6 (2019).

32 Hubert, L. & Arabie, P. Comparing partitions. J Classif 2, 193–218, doi:10.1007/bf01908075 (1985).

33 Rousseeuw, P. J. Silhouettes: A graphical aid to the interpretation and validation of cluster analysis. J Comput Appl Math 20, 53–65, doi:10.1016/0377-0427(87)90125-7 (1987).

34 Grubman, A. et al. A single-cell atlas of entorhinal cortex from individuals with Alzheimer’s disease reveals cell-type-specific gene expression regulation. Nature Neuroscience 22, 2087–2097, doi:10.1038/s41593-019-0539-4 (2019).

35 Tanzi, R. E. The genetics of Alzheimer disease. Cold Spring Harb Perspect Med 2, a006296, doi:10.1101/cshperspect.a006296 (2012).

36 Su, B. et al. Oxidative stress signaling in Alzheimer’s disease. Curr Alzheimer Res 5, 525–532, doi:10.2174/156720508786898451 (2008).

37 Ma, A. et al. IRIS3: integrated cell-type-specific regulon inference server from single-cell RNA-Seq. Nucleic Acids Research, doi:10.1093/nar/gkaa394 (2020).

38 Karch, C. M., Ezerskiy, L. A., Bertelsen, S., Alzheimer’s Disease Genetics, C. & Goate, A. M. Alzheimer’s Disease Risk Polymorphisms Regulate Gene Expression in the ZCWPW1 and the CELF1 Loci. PloS one 11, e0148717’e0148717, doi:10.1371/journal.pone.0148717 (2016).

39 Franzén, O., Gan, L.-M. & Björkegren, J. L. M. PanglaoDB: a web server for exploration of mouse and human single-cell RNA sequencing data. Database 2019, doi:10.1093/database/baz046 (2019).

40 Mathys, H. et al. Single-cell transcriptomic analysis of Alzheimer’s disease. Nature 570, 332–337, doi:10.1038/s41586-019-1195-2 (2019).

41 Yamakawa, H. et al. The Transcription Factor Sp3 Cooperates with HDAC2 to Regulate Synaptic Function and Plasticity in Neurons. Cell Rep 20, 1319–1334, doi:10.1016/j.celrep.2017.07.044 (2017).

42 Boutillier, S. et al. Sp3 and sp4 transcription factor levels are increased in brains of patients with Alzheimer’s disease. Neuro-degenerative diseases 4, 413–423, doi:10.1159/000107701 (2007).

43 Hu, Z., Dong, Y., Wang, K. & Sun, Y. Heterogeneous Graph Transformer. Proceedings of The Web Conference 2020 (2020).

44 Ma, A., McDermaid, A., Xu, J., Chang, Y. & Ma, Q. Integrative Methods and Practical Challenges for Single-Cell Multi-omics. Trends in Biotechnology, doi:10.1016/j.tibtech.2020.02.013 (2020).

45 Han, A., Glanville, J., Hansmann, L. & Davis, M. M. Linking T-cell receptor sequence to functional phenotype at the single-cell level. Nature Biotechnology 32, 684–692, doi:10.1038/nbt.2938 (2014).

46 Grün, D. Revealing dynamics of gene expression variability in cell state space. Nat Methods 17, 45–49, doi:10.1038/s41592-019-0632-3 (2020).

47 Liu, F. T., Ting, K. M. & Zhou, Z. in 2008 Eighth IEEE International Conference on Data Mining. 413–422.

48 Lin, P., Troup, M. & Ho, J. W. J. G. b. CIDR: Ultrafast and accurate clustering through imputation for single-cell RNA-seq data. 18, 59 (2017).

49 Qiu, X. et al. Reversed graph embedding resolves complex single-cell trajectories. Nat Methods 14, 979–982, doi:10.1038/nmeth.4402 (2017).

50 Lin, P., Troup, M. & Ho, J. W. CIDR: Ultrafast and accurate clustering through imputation for single-cell RNA-seq data. Genome Biol 18, 59, doi:10.1186/s13059-017-1188-0 (2017).

51 Subramanian, A. et al. Gene set enrichment analysis: a knowledge-based approach for interpreting genome-wide expression profiles. Proc Natl Acad Sci U S A 102, 15545–15550, doi:10.1073/pnas.0506580102 (2005).

