## Supplementary Figure S1-S7, Table S1-S2, Methods S1-S4, Algorithm S1 for "scGNN: a novel graph neural network framework for single-cell RNA-Seq analyses"

**Supplementary Figures**

Figure S1. Workflow of the iterative process in scGNN.

Figure S2. An illustration of LTMG modeling.

Figure S3. Comparison of gene co-expression relationships in the Klein dataset.

Figure S4. Design of ablation tests and parameter searching of scGNN.

Figure S5. Clustering results of scGNN compared to existing clustering tools.

Figure S6. Comparison of DEG expression before and after scGNN imputation.

Figure S7. A full list of enriched pathways using DEGs between AD and control cells within each cluster.

**Supplementary Tables**

Table S1. Description of datasets and third-party tools for performance comparisons.

Table S2: Design of ablation tests in clustering.

Table S3: Ablation tests with or without graph embedding

Table S4: Comparison between naïve PCA and PCA with graph embedding

Table S5: Comparison between scGNN and PCA with graph embedding

Table S6: Ablation tests with regulatory signal integration

Table S7: Ablation tests with cluster autoencoders

Table S8: Choosing K and intensities in clustering on the Klein Dataset

Table S9: Parameter searching in imputation on the Klein Dataset

Table S10: Parameter searching in imputation on the Zeisel Dataset

Table S11: Ablation tests in imputation

Table S12: Parameter searching in L1/L2 terms in the imputation

Table S13: CTSRs predicted from IRIS3 using the scGNN imputed matrix.

Table S14: SP3 regulated genes in OPC, Astrocyte, and Neuron clusters.

**Supplementary Methods**

Algorithm 1. Pseudo code of scGNN

Method S1. Ablation tests on computational components in scGNN

Method S2. Robustness of scGNN to hyperparameter choices

Method S3. Parameter tuning and ablation tests in imputation tasks in scGNN

Method S4. Quantitative criteria used in the study

**Supplementary Figures**

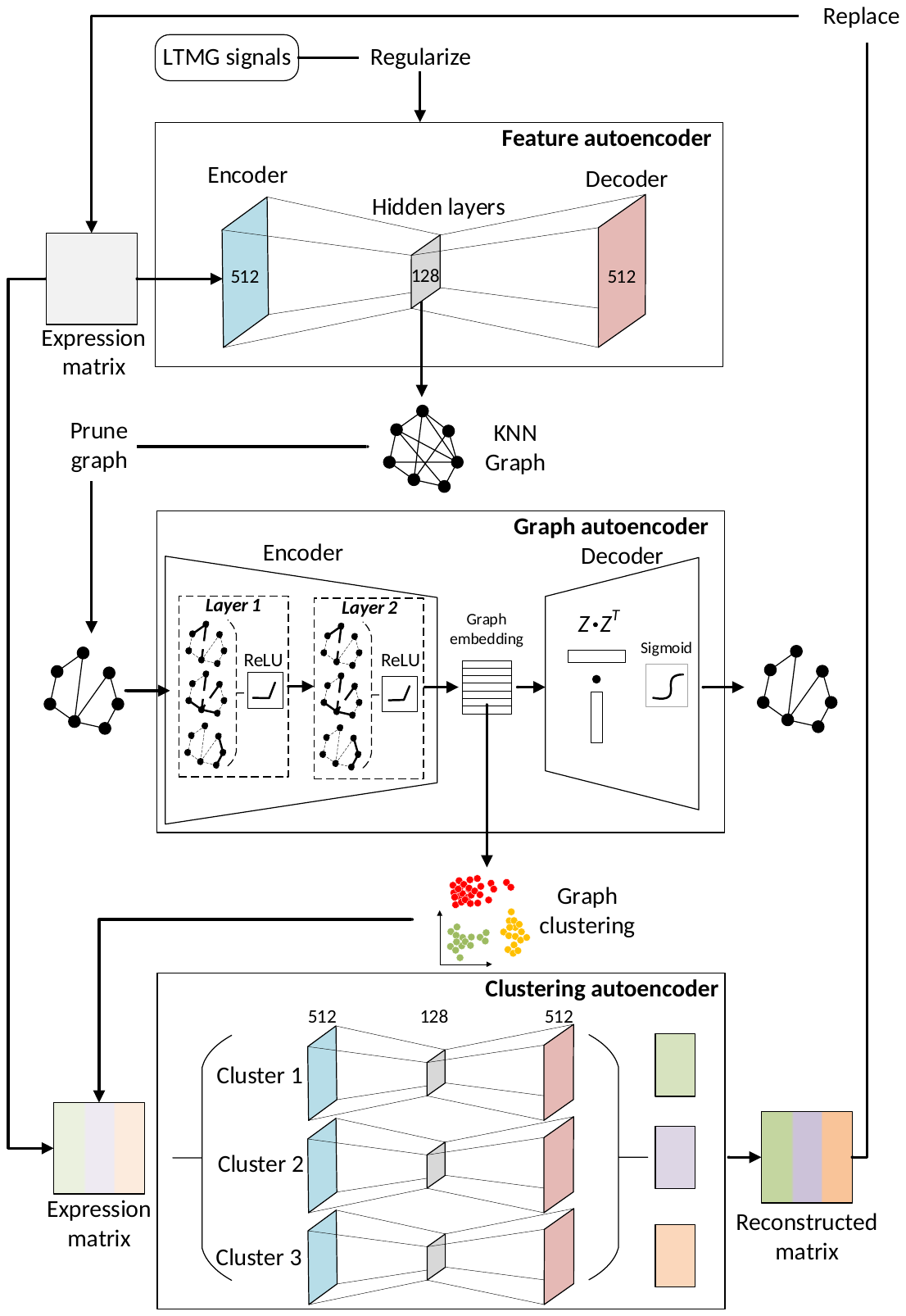

**Figure S1**. Workflow of the iterative process in scGNN.

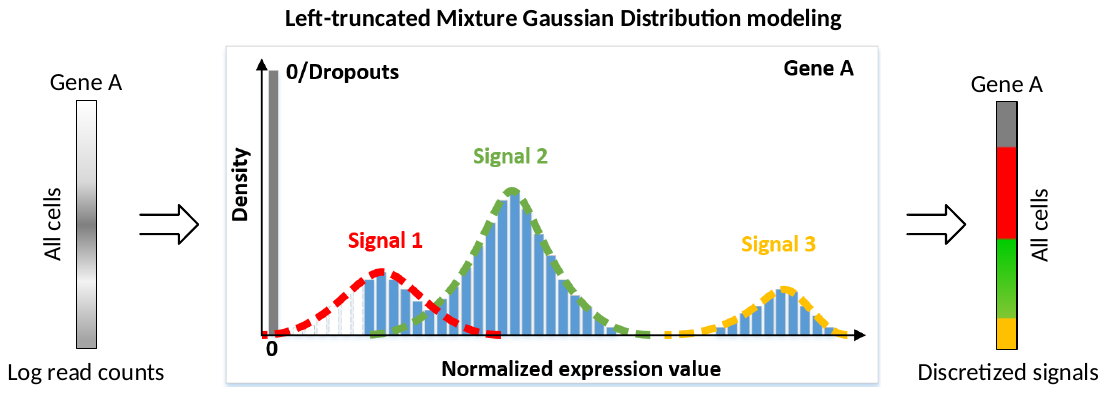

**Figure S2**. An illustration of LTMG modeling. For Gene A with log read counts in the raw scRNA-Seq matrix, LTMG models the expression values into multiple Gaussian distributions to indicate its underlying regulatory signals. Expression values in cells within the same distribution will be discretized into the same number. For instance, expressions in cells with signal 1 (red), 2 (green), and 3 (yellow) are transformed into 1, 2, and 3, respectively. All zero values will remain the same.

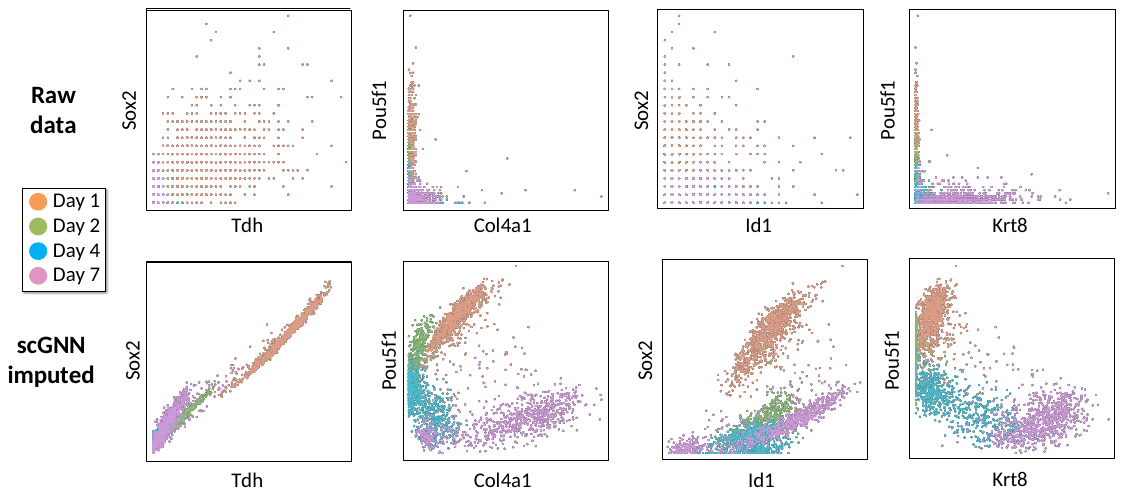

**Figure S3**. Comparison of gene co-expression relationships in the Klein dataset. Different colors indicate cell clusters given in the original paper (Day 1, 2, 7, and 9).

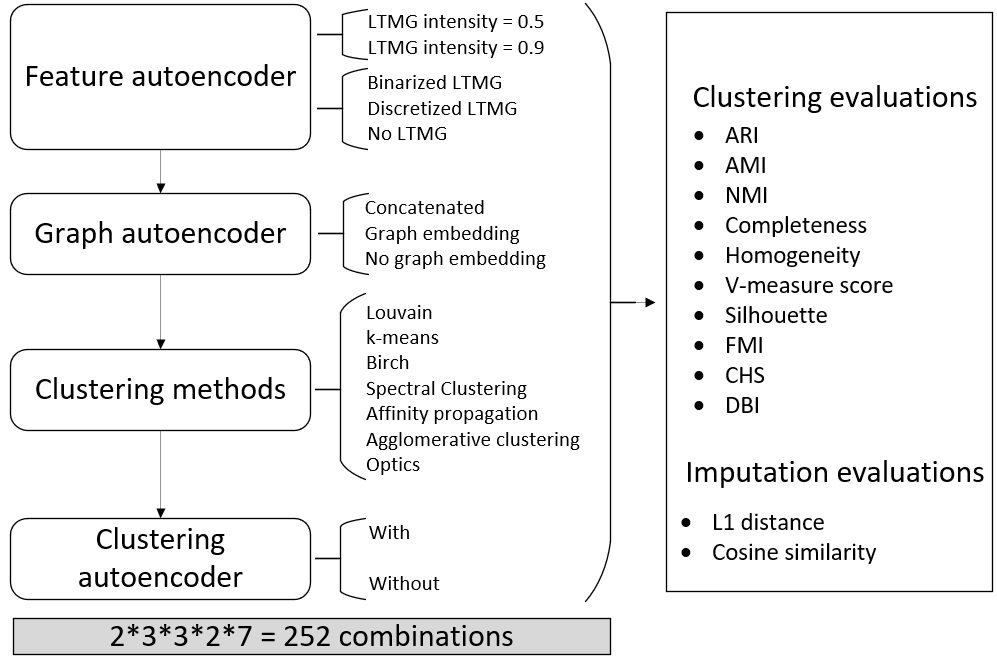

**Figure S4**. Design of ablation tests and parameter searching of scGNN. We tested 252 parameter combinations in scGNN for the three autoencoders in the iterative process to select the best scGNN performance. All results are evaluated via ten criteria for clustering and two for imputation.

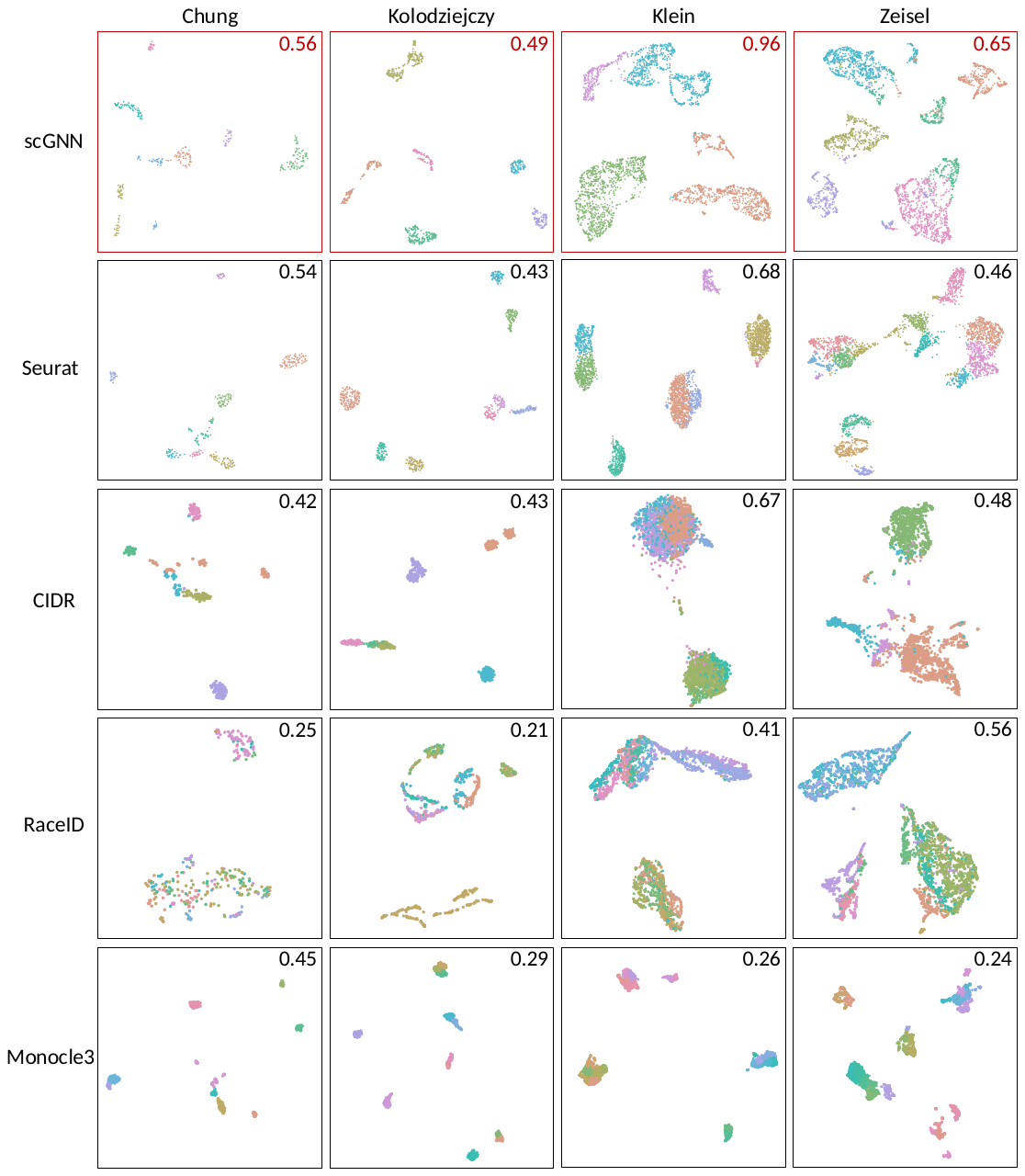

**Figure S5**. Clustering results of scGNN compared to existing clustering tools. The comparison was conducted on four tools (i.e., Seurat, CIDR, RaceID, and Monocle3) using four benchmark datasets. ARI of each test is indicated on each UMAP, comparing the predicted cell clusters to the benchmark labels.

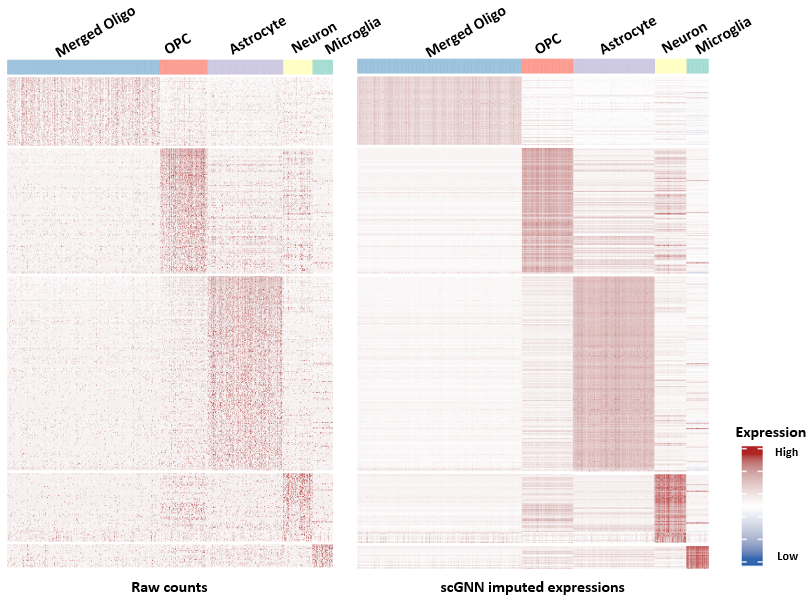

**Figure S6**. Comparison of DEG expression before (Left) and after scGNN imputation (Right). DEGs were identified using the Seurat package based on scGNN predicted clusters, and the six oligodendrocyte sub-clusters were merged into one. Cells were randomly selected from half of the merged oligo group to make the figure more balanced.

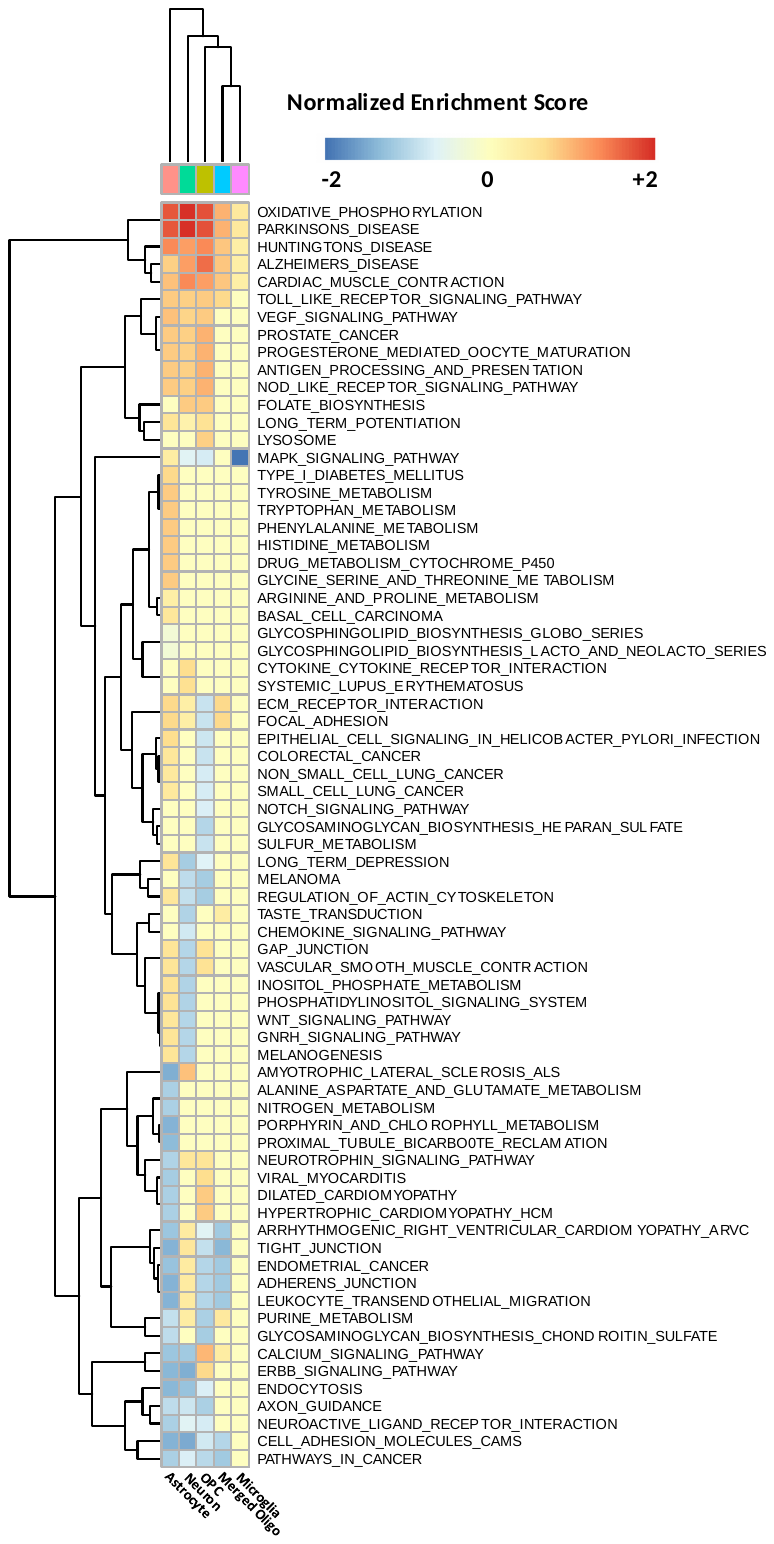

**Figure S7**. A full list of enriched pathways using DEGs between AD and control cells within each cluster. The enrichment test was conducted by using GSEA. Colors indicate normalized enrichment scores.

**Supplementary Tables**

**Table S1**. Description of datasets and third-party tools for performance comparisons.

| **Datasets used in paper** | | | | | |
| --- | --- | --- | --- | --- | --- |
| **Dataset** | **# of genes** | **# of cells** | **# clusters** | **Data index** | **Ref** |
| Klein’s | 24,175 | 2,717 | 4 | GSE65525 | ^1^ |
| Zeisel’s | 19,972 | 3,005 | 9 | GSE60361 | ^2^ |
| Chung | 57,915 | 317 | 4 | GSE75688 | ^3^ |
| Kolodziejczy | 38,653 | 704 | 3 | E-MTAB-2600 | ^4^ |
| AD data | - | 13,214 | 8 | GSE138852 | ^5^ |
| **Tools for performance comparison** | | | | | |
|  | **Tool name** | **Version** | **Implementation** | **Parameters** | **Ref** |
| Imputation tools | MAGIC | 2.0 | R & Python | Default | ^6^ |
|  | SAUCIE | - | Python |  | ^7^ |
|  | SAVER | 1.1.2 | R |  | ^8^ |
|  | scImpute | 0.0.9 | R |  | ^9^ |
|  | scVI | 0.6.5 | Python |  | ^10^ |
|  | DCA | - | Python |  | ^11^ |
|  | DeepImpute | 1.1 | Python |  | ^12^ |
| Clustering tools | Seurat | 3.1 | R |  | ^13^ |
|  | CIDR | 0.1.5 | R & C++ |  | ^14^ |
|  | Monocle | 3.0 | R |  | ^15^ |
|  | RaceID | - | R |  | ^16^ |

**Table S2**: Design of ablation tests in clustering. Test numbers are organized for different combinations of computational components.

| Test NO. | Clustering  Method | With/Without Cluster Autoencoder | Regulation Types Selection | Embedding Types  Selection |
| --- | --- | --- | --- | --- |
| 1 | Louvain/  K-Means/  Birch/  Spectral clustering/ Agglomerative clustering/  Affinity propagation/ Optics | Without | Discretized | Concatenated |
| 2 |  |  |  | Graph Embedding |
| 3 |  |  |  | No Graph Embedding |
| 4 |  |  | Binarized | Concatenated |
| 5 |  |  |  | Graph Embedding |
| 6 |  |  |  | No Graph Embedding |
| 7 |  |  | No | Concatenated |
| 8 |  |  |  | Graph Embedding |
| 9 |  |  |  | No Graph Embedding |
| 10 |  | With | Discretized | Concatenated |
| 11 |  |  |  | Graph Embedding |
| 12 |  |  |  | No Graph Embedding |
| 13 |  |  | Binarized | Concatenated |
| 14 |  |  |  | Graph Embedding |
| 15 |  |  |  | No Graph Embedding |
| 16 |  |  | No | Concatenated |
| 17 |  |  |  | Graph Embedding |
| 18 |  |  |  | No Graph Embedding |

**Supplemental Methods**

**Algorithm S1. Pseudo code of scGNN**

**Algorithm 1:** scGNN

**Input:** scRNA-seq expression $X_{raw}$

**Output:** Clustering results $R$, cell graph $A$, imputed expression $\hat{X}$

$TRS$ 🡨 Left-Truncated-Mixed-Gaussian Model ($X_{raw}$)

Embedding $X$’, reconstructed $X$ 🡨 Feature Autoencoder ($X_{raw}$,$TRS$)

$A$ 🡨 Build KNN Graph ($X$’)

Graph Embedding $Z$ 🡨 Graph Autoencoder ($A,X$’)

$R$ 🡨 Clustering ($Z$)

$X$ 🡨 Cluster Autoencoder ($X,R$)

**While** $R$ is not converge or $X$ is not converge **do:**

Embedding $X$’, updated $X$ 🡨 Feature Autoencoder ($X$)

$A$ 🡨 Build KNN Graph($X$’)

Graph Embedding $Z$ 🡨 Graph Autoencoder ($A,X$’)

$R$ 🡨 Clustering ($Z$)

$X$ 🡨 Cluster Autoencoder ($X,R$)

**End**

Imputed expression $\hat{X}$ 🡨 Imputation Autoencoder($X_{raw},R,A$)

**Return** $R$,$A$,$\hat{X}$

**Method S1. Ablation tests on computational components in scGNN**

scGNN comprises several comprehensive computational components to infer cell clusters and impute gene expressions. In this section, we explore the essential roles of scGNN components and the concepts of ablation tests in the benchmark experiments.

The explored components in scGNN are listed as (a) *embedding selection* from clustering on feature autoencoder embedding, graph embedding, or feature autoencoder embedding concatenated with graph embedding; (b) choosing whether or not to use *integrating regulatory signals* in the feature autoencoder; (c) weak/strong intensities of *regulatory regularization intensities*; (d) binarized/discretized *regulatory regularization selection*; (e) deciding whether to use an *autoencoder*; (f) *clustering methods selection* from Louvain^17^, k-means^18^, Birch^19^, spectral clustering^20^, agglomerative clustering^21^, affinity propagation^22^, and Optics^23^.

The goal of the ablation test is to design several scenarios by only excluding one of these computational components; then, check its influences in the clustering performance changes quantitatively. If the performance gets worse, it means the component tested contributes to scGNN. Here are the details of the components:

(a) Embedding selection. Given the single-cell graph, graph embedding integrates the topological information of the cell-cell relationships using graph autoencoders by aggregating the node neighbors with graph convolution networks. In scGNN, the single-cell graph is built from the preceding feature autoencoder embedding. Since learned graph embedding is the main contribution of the proposed scGNN, scGNN identifies cell types by clustering on graph embeddings. To validate the efficiency of graph embedding, we also tested the performances without/with graph embeddings, i.e. clustering on the embedding learned from the main feature autoencoder, or only using the input of the graph autoencoder. We also tested another possible solution, concatenating the embedding from the main feature autoencoder and the graph embedding from the graph autoencoder together.
(b) Deciding if or when to integrate regulatory signals. Integrating regulatory signals from scRNA expression through regularizing LTMG in the feature autoencoder is another main contribution of scGNN. To validate the efficiency of integrating regulatory signals in scGNN, we tested both scenarios with/without the regularization terms (*Eq*.5 in the main text) in the loss function of the feature autoencoder.
(c) Intensity of regularization. When integrating regulatory signals in the scGNN model, it may also be essential to check the appropriate intensity of the regularization. As shown in Eq. 5 and Eq. 6, $\alpha$ is set at different intensities of 0.5 and 0.9 as the weak and strong intensities, respectively. In other words, the penalty intensities of genes with different regulatory roles have different impacts on the loss function training. For example, when $\alpha$ is set to 0.5, the impact of a gene with an LTMG value 1 is twice that of the gene with an LTMG value of 0 in the loss function. When $\alpha$ is set to 0.9, the impact is 10 times greater than that of a gene with an LTMG value of 1 and a gene with an LTMG value of 0.
(d) Regulation selection. The output of regulatory signals modeled by LTMG (TRS in *Eq*.5 and *Eq*.6) may contain two types of values: One value is discretized regulatory signals given as discrete numbers such as 0/1/2/3 in the main text; the other value is binary signals given as 0/1, which concludes all the non-zero values to 1. These values are passed directly to the feature autoencoder and results in different types of penalties. We named the former as discretized regularization and the latter as binarized regularization.
(e) Whether using cluster autoencoders. In each iteration of scGNN, each cluster autoencoder corresponds to each identified cluster autoencoder. In practical usage, cluster autoencoders may consume extensive computational time if too many Imputations are identified. We tested the performances with and without the usage of cluster autoencoders. If not using cluster autoencoders, the reconstructed expression from the feature autoencoder is directly passed to the next iteration.
(f) Clustering method selection. Several well-established clustering methods such as Louvain, K-Means, Birch, spectral clustering, agglomerative clustering, affinity propagation, and Optics were tested in the framework of scGNN. Here, we only tested the performances on the learned embedding of the last iteration in scGNN, which was used as the final clustering output of scGNN. All these clustering methods were tested individually for each test. K-Means, Birch, spectral clustering, agglomerative clustering, affinity propagation, and Optics used python implementations of scikit-learn (version 0.22.1). Louvain used the R package igraph (version 1.2.4.2). We used the default parameters of these methods in our tests. For algorithms that require the number of clusters as the input, such as K-Means and Birch, we used Louvain first and took the number of clusters identified by Louvain for those algorithms.

The embedding selection, regulation integration, imputation autoencoders, and iteration processes are independent from each other and contribute differently to the performances of scGNN. The ablation tests are designed in the following combinations of the computational components in **Figure S5** and **Table S2**. All the tests are tested quantitatively on the four benchmark datasets.

To test the capacity of graph embedding, we also added two baseline tests using PCA to replace the feature autoencoder:

- PCA without graph embedding: Using PCA to get 100 principal components from the input scRNA expression and clustering on these PCA results.
- PCA graph embedding: Using PCA to get 100 principal components from the input scRNA expression, building a single-cell graph using these PCs, and clustering on the learned graph, embedding from the graph autoencoder.

From these ablation tests results, we can see:

1. Birch and k-means are consistently better than other clustering methods. From all the tests, each of the learned embeddings in different benchmarks is clustered by different clustering methods. Comparing with other candidate clustering methods, Birch and K-Means are consistently better or as good as the best clustering methods. We chose K-Means as the default clustering method in scGNN. In the following tests, we mainly explore the performances of K-Means.
2. scGNN with graph embedding is better than without embedding. After comparing all the possible conditions with/without graph embedding, we checked results with discretized regularization without/with cluster autoencoder, binarized regularization without/with cluster autoencoder, and no regulatory signal integration without/with cluster autoencoder. All the results are detailed in **Table S3**. Most of the results demonstrate that scGNN in graph embedding has higher ARI and silhouette scores than clustering on the feature autoencoder embeddings. In the meantime, the concatenated embedding does not bring better results than solely using graph embedding.
3. Graph embedding works better than the baseline naïve PCA. As single-cell graphs are built for the graph autoencoder in scGNN, we also checked whether graph embedding from graph autoencoder improves performances by building single-cell graphs from other dimension deduction methods such as PCA. **Table S4** compares naïve PCA with graph embedding PCA. Adding graph embedding significantly improves the clustering results in most benchmarks, especially in datasets with more cells. These results indicate graph embedding itself increases cell type clustering performance.
4. scGNN is better than graph embedding PCA. Compared with graph embedding PCA, scGNN not only adopts graph embedding but also involves other innovations such as regulatory signals integration and iterative processing. With all tests with graph embedding with/without cluster autoencoder along with discretized/binarized/no-signals, **Table S5** compares these conditions with graph embedding PCA. Results show that scGNN works better in most benchmarks.
5. scGNN with integrating regulatory signals provides better clustering results. After comparing each of the three possible yet different conditions in regulatory signals integration with discretized regularization, with binarized regularization and no integration, we tested without/with graph embedding along with a without/with cluster autoencoder. **Table S6** details the compared results. scGNN has better results with regulatory signals integration, regardless of either discretized regulation or binarized regulation.
6. The strong intensity of regulation regularization is better than the weak intensity of regularization. Two different intensities of regulation regularizations are compared with only change in the $\alpha$ parameter in *Eq*.(5). Although the best choice of $\alpha$ may depend on the data, the strong intensity ($\alpha=0.9$) of regulation regularization has better performances than the weak regularization intensity ($\alpha=0.5$) in both discretized and binarized signals integrations. **Table S6** details the comparisons.
7. The cluster autoencoder brings better results to scGNN. Comparing pairwise tests with/without the cluster autoencoder, all possible tests in discretized, binarized, and non-regulatory signal integration with/without graph embedding are shown in **Table S7**. The cluster autoencoder clearly brought better results to scGNN.

From these observations of the ablation tests, graph embedding, regulation integration, cluster autoencoder, and iterative processes, we found that all these components worked together in scGNN to contribute to the cell type clustering accuracy.

**Method S2. Robustness of scGNN to hyperparameter choices**

We used the quality criteria described above to address the robustness of scGNN to hyper-parameter choices for a different *K* in the stage of building a cell graph using K-Nearest-Neighbor and LTMG intensities. **Table S8** demonstrates the clustering results for the dataset Klein. *K* is selected as 5, 10, 20, 30, and 40. In each K, the LTMG intensity $\alpha$ was selected as the gradient of 0.1, 0.25, 0.5, 0.75 and 0.9. Stronger LTMG intensity usually leads to better results if the choice of K is not too small. These results demonstrate that scGNN is robust in selecting hyperparameters and that it does not significantly affect clustering quality.

**Method S3. Parameter tuning and ablation tests in scGNN imputation tasks**

We used a similar grid search parameter tuning strategy in imputation tasks after getting clustering results from the iteration. The imputation autoencoder adopted *Eq*. 14 in the main text to impute. Using the gradient in 0.0/0.1/0.3/0.9, the LTMG intensity $\alpha$, graph intensity $\gamma_{1},$ and cell type intensity $\gamma_{2}$ were tested comprehensively in 10% of the drop events in the benchmark Klein dataset as shown in **Table S9** and in the Zeisel dataset in **Table S10**. Considering random effects from flipping nonzero to zeros brought from different seeds, each dataset repeats three times with seeds 1,2,3, which we examined as the average and standard deviation. The mean, median, min, and max L1 errors between the imputed and original expression were used as the quantitative measurements. We agreed that the imputation results were robust and insensitive to variation in most of these intensities.

We performed ablation tests in imputation for the best parameter set 0.0/0.3/0.1 with/without graph autoencoder, LTMG integration, and cluster autoencoder in the iterative stages of clustering results. **Table S10** shows the results. The mean, median, min, and max L1 errors between the imputed and original expression are used as the quantitative measurements. Even though the results were not perfectly consistent with clustering ablation test results, these imputation results demonstrate the efficiency of these computational components in regularizing imputation results.

We also tested the effectiveness of involving L1 or L2 terms in the loss of function in the imputation autoencoder. Using the intensity parameter set in **Table S12** shows tests in the different permutation of L1/L2 intensities of 0.001/0.01/0.1/1 of scGNN. From the quantitative measurements in the mean, median, min, max L1 errors, and cosine similarity, we found that only using the L1 term gets better results in imputation.

**Method S4. Quantitative criteria used in the study**

Besides the ARI and average silhouette scores introduced in the main text, we also adopted adjusted and normalized mutual information^24^, completeness^25^, homogeneity^25^, V-measure score^25^, the Fowlkes-Mallows index^26^, the Calinski-Harabasz score^27^, and the Davies-Boudin index^28^ to compute similarities between inferred clustering results and gold-standard benchmarks.

**AMI** (Adjusted mutual information) is an adjustment of the mutual information (MI) score to account for chance. $X$ and $Y$ are the inferred clustering results and the benchmark clustering results. Higher $AMI\in[0,1]$ means higher similarity

$$AMI(X,Y)=\frac{MI(X,Y)-E(MI(X,Y))}{Avg(H(X,Y))-E(MI(X,Y))}$$

Entropy $H$ is the amount of uncertainty for a partition set, defined by:

$$H(X)=-\sum_{i=1}^{|X|} P(i)\log(P(i))$$

where $P(i)=\left| X_{i} \right|/N$. The mutual information (MI) between $X$ and $Y$ is calculated by:

$$MI(X,Y)=\sum_{i=1}^{|X|} \sum_{j=1}^{|Y|} P(i,j)\log(\frac{P(i,j)}{P(i)P(j)})$$

The expectation of MI can be calculated from:

$$E\left[ MI(X,Y)) \right]=\sum_{i=1}^{|X|} \sum_{j=1}^{|Y|} \sum_{n_{i,j}=(a_{i}+b_{j}-N)}^{\min(a_{i},b_{j})} \frac{n_{ij}}{N}\log(\frac{N\cdot n_{ij}}{a_{i}b_{j}})\frac{a_{i}!b_{j}!(N-a_{i})!(N-b_{j})!}{N!n_{ij}!(a_{i}-n_{ij})!(b_{j}-n_{ij})!(N-a_{i}-b_{j}+n_{ij})!}$$

where $a_{i}$is the number of elements in $X_{i}, andb_{i}$ is the number of elements in $Y_{i}$.

**NMI** (Normalized mutual information) is another adjusted form of mutual information (MI). It is defined as:

$$NMI(X,Y)=\frac{MI(X,Y)}{mean(H(X),H(Y))}$$

$\mathrm{Higher}AMI\in[0,1]$ means higher similarity.

**Homogeneity** measures to what extent each cluster contains only members of a single class. Higher $Homogeneity\in[0,1]$ means higher similarity.

$$homogeneity=1-\frac{H(C|K)}{H(C)}$$

where $H(C|K)$ is the conditional entropy of the classes given as the cluster assignments and are written as:

$$H(C|K)=-\sum_{c=1}^{|C|} \sum_{k=1}^{|K|} \frac{n_{c,k}}{n}\cdot\log(\frac{n_{c,k}}{n_{k}})$$

thus, $H(C)$ is the entropy of the classes defined as:

$$H(C|K)=-\sum_{c=1}^{|C|} \frac{n_{c}}{n}\cdot\log(\frac{n_{c}}{n})$$

where $n$ is the total number of samples. $n_{c}$and$n_{k}$ are the number of samples, belonging to class $c$and cluster $k$, respectively. $n_{c,k}$is the number of samples from class $c$assigned to cluster $k$.

**Completeness** measures how much all members of a given class are assigned to the same cluster. Higher $Completeness\in[0,1]$ means higher similarity.

$$completeness=1-\frac{H(K|C)}{H(K)}$$

**V-measure** is the harmonic mean of homogeneity and the completeness score.

$$Vmeasure=\frac{2\times\hom ogeneity\times completeness}{\hom ogeneity+completeness}$$

$\mathrm{Higher}Vmeasure\in[0,1]$ means higher similarity.

**Fowlkes-Mallows index** (FMI) is defined as the geometric means of the pairwise precision and recall:

$$FMI=\frac{TP}{\sqrt{(TP+FP)(TP+FN)}}$$

where TP is the true positive, FP is the false positive, and FN is the false negative.

$A higher FMI\in[0,1]$ means a higher similarity.

**Calinski-Harabasz score**, also known as the variance ratio criterion, is the ratio of the between-clusters dispersion and inter-cluster dispersion for all clusters, where dispersion is defined as the sum of distances squared. For a set of data $E$ of size $n_{E}$ which is clustered into $k$ clusters, the Calinski-Harabasz score *s* is defined as the ratio of the between-clusters dispersion mean and the within-cluster dispersion:

$$s=\frac{tr(B_{k})}{tr(W_{k})}\times\frac{n_{E}-k}{k-1}$$

where $tr(B_{k})$ is the trace of the between-group dispersion matrix and $tr\left( W_{k} \right)$is the trace of the within-cluster dispersion matrix defined by:

$$W_{k}=\sum_{q=1}^{k} \sum_{x\in C_{q}} (x-c_{q})(x-c_{q})^{T}$$

$$B_{k}=\sum_{q=1}^{k} n_{q}(c_{q}-c_{E})(c_{q}-c_{E})^{T}$$

where $C_{q}$is the set of points in cluster $q$;$c_{q}$is the center of cluster $q$; $C_{E} \mathrm{is}$the center of $E$, and $n_{q}$is the number of points in cluster $q$.

**Davies-Boudin index** signifies the average similarity between clusters, where the similarity is a measure that compares the distance between clusters with the size of the clusters themselves. A lower value indicates a better partition.

$$DaviesBouldin=\frac{1}{k}\sum_{i=1}^{k} \max_{i\neq j}\frac{s_{i}+s_{j}}{d_{ij}}$$

where $s_{i}$is the average distance between each point of cluster $i$, and the centroid of that cluster, also known as the cluster diameter. $d_{ij}$is the distance between cluster centroids $i$and $j$.

**References**

1	Klein, A. M. *et al.* Droplet barcoding for single-cell transcriptomics applied to embryonic stem cells. *Cell* **161**, 1187-1201, doi:10.1016/j.cell.2015.04.044 (2015).

2	Zeisel, A. *et al.* Brain structure. Cell types in the mouse cortex and hippocampus revealed by single-cell RNA-seq. *Science* **347**, 1138-1142, doi:10.1126/science.aaa1934 (2015).

3	Chung, W. *et al.* Single-cell RNA-seq enables comprehensive tumour and immune cell profiling in primary breast cancer. *Nat Commun* **8**, 15081, doi:10.1038/ncomms15081 (2017).

4	Kolodziejczyk, A. A. *et al.* Single Cell RNA-Sequencing of Pluripotent States Unlocks Modular Transcriptional Variation. *Cell Stem Cell* **17**, 471-485, doi:10.1016/j.stem.2015.09.011 (2015).

5	Grubman, A. *et al.* A single-cell atlas of entorhinal cortex from individuals with Alzheimer's disease reveals cell-type-specific gene expression regulation. *Nat Neurosci* **22**, 2087-2097, doi:10.1038/s41593-019-0539-4 (2019).

6	van Dijk, D. *et al.* Recovering Gene Interactions from Single-Cell Data Using Data Diffusion. *Cell* **174**, 716-729 e727, doi:10.1016/j.cell.2018.05.061 (2018).

7	Amodio, M. *et al.* Exploring single-cell data with deep multitasking neural networks. *Nat Methods* **16**, 1139-1145, doi:10.1038/s41592-019-0576-7 (2019).

8	Huang, M. *et al.* SAVER: gene expression recovery for single-cell RNA sequencing. *Nat Methods* **15**, 539-542, doi:10.1038/s41592-018-0033-z (2018).

9	Li, W. V. & Li, J. J. An accurate and robust imputation method scImpute for single-cell RNA-seq data. *Nat Commun* **9**, 997, doi:10.1038/s41467-018-03405-7 (2018).

10	Lopez, R., Regier, J., Cole, M. B., Jordan, M. I. & Yosef, N. Deep generative modeling for single-cell transcriptomics. *Nat Methods* **15**, 1053-1058, doi:10.1038/s41592-018-0229-2 (2018).

11	Eraslan, G., Simon, L. M., Mircea, M., Mueller, N. S. & Theis, F. J. Single-cell RNA-seq denoising using a deep count autoencoder. *Nat Commun* **10**, 390, doi:10.1038/s41467-018-07931-2 (2019).

12	Arisdakessian, C., Poirion, O., Yunits, B., Zhu, X. & Garmire, L. X. DeepImpute: an accurate, fast, and scalable deep neural network method to impute single-cell RNA-seq data. *Genome Biol* **20**, 211, doi:10.1186/s13059-019-1837-6 (2019).

13	Butler, A., Hoffman, P., Smibert, P., Papalexi, E. & Satija, R. Integrating single-cell transcriptomic data across different conditions, technologies, and species. *Nat Biotechnol* **36**, 411-420, doi:10.1038/nbt.4096 (2018).

14	Lin, P., Troup, M. & Ho, J. W. J. G. b. CIDR: Ultrafast and accurate clustering through imputation for single-cell RNA-seq data. **18**, 59 (2017).

15	Qiu, X. *et al.* Reversed graph embedding resolves complex single-cell trajectories. *Nat Methods* **14**, 979-982, doi:10.1038/nmeth.4402 (2017).

16	Lin, P., Troup, M. & Ho, J. W. CIDR: Ultrafast and accurate clustering through imputation for single-cell RNA-seq data. *Genome Biol* **18**, 59, doi:10.1186/s13059-017-1188-0 (2017).

17	Blondel, V. D., Guillaume, J.-L., Lambiotte, R. & Lefebvre, E. Fast unfolding of communities in large networks. *Journal of Statistical Mechanics: Theory and Experiment* **2008**, P10008, doi:10.1088/1742-5468/2008/10/p10008 (2008).

18	Hartigan, J. A. & Wong, M. A. Algorithm AS 136: A K-Means Clustering Algorithm. *Journal of the Royal Statistical Society. Series C (Applied Statistics)* **28**, 100-108, doi:10.2307/2346830 (1979).

19	Zhang, T., Ramakrishnan, R. & Livny, M. BIRCH: an efficient data clustering method for very large databases. **25**, 103–114, doi:10.1145/235968.233324 (1996).

20	von Luxburg, U. A tutorial on spectral clustering. *Statistics and Computing* **17**, 395-416, doi:10.1007/s11222-007-9033-z (2007).

21	Beeferman, D. & Berger, A. in *Proceedings of the sixth ACM SIGKDD international conference on Knowledge discovery and data mining*	407–416 (Association for Computing Machinery, Boston, Massachusetts, USA, 2000).

22	Frey, B. J. & Dueck, D. Clustering by Passing Messages Between Data Points. **315**, 972-976, doi:10.1126/science.1136800 %J Science (2007).

23	Ankerst, M., Breunig, M. M., Kriegel, H.-P. & Sander, J. OPTICS: ordering points to identify the clustering structure. **28**, 49–60, doi:10.1145/304181.304187 (1999).

24	Vinh, N. X., Epps, J. & Bailey, J. Information Theoretic Measures for Clusterings Comparison: Variants, Properties, Normalization and Correction for Chance. **11**, 2837–2854 (2010).

25	Rosenberg, A. & Hirschberg, J. in *Proceedings of the 2007 joint conference on empirical methods in natural language processing and computational natural language learning (EMNLP-CoNLL).* 410-420.

26	Fowlkes, E. B. & Mallows, C. L. A Method for Comparing Two Hierarchical Clusterings. *Journal of the American Statistical Association* **78**, 553-569, doi:10.1080/01621459.1983.10478008 (1983).

27	Calinski, T. & Harabasz, J. A dendrite method for cluster analysis. *Communications in Statistics - Theory and Methods* **3**, 1-27, doi:10.1080/03610927408827101 (1974).

28	Halkidi, M., Batistakis, Y. & Vazirgiannis, M. On Clustering Validation Techniques. *Journal of Intelligent Information Systems* **17**, 107-145, doi:10.1023/A:1012801612483 (2001).
